# Zebrafish *Danio rerio* trunk muscle structure and growth from a spatial transcriptomics perspective

**DOI:** 10.1101/2021.06.03.446321

**Authors:** Guanting Liu, Takumi Ito, Yusuke Kijima, Kazutoshi Yoshitake, Shuichi Asakawa, Shugo Watabe, Shigeharu Kinoshita

## Abstract

Compared to mammals, some fish exhibit indeterminate growth characteristics, meaning they can continue growing throughout their lives. Zebrafish trunk skeletal muscle can in general be classified into slow, intermediate, and fast based on morphological and physiological characteristics. After hatching, hyperplasia can be observed in the muscles of juvenile zebrafish, and with growth, hyperplasia in the fast muscles gradually decreases until it stagnates, after which fast muscle development relies on hypertrophy. In slow muscle, hyperplasia continues throughout life. Teleost muscle structure and growth has been described mainly by morphological and physiological features based on the expression of a limited number of proteins, transcripts, and metabolites. The details of mechanism remain unclear. Visium Spatial Gene Expression solution was used in this study. On the adult slide, 10 clusters were obtained based on whole gene expression similarities. The spatial expression of myosin heave chains, myosin light chains and myosin-binding proteins was investigated. GO enrichment analysis was also performed on different muscle regions of aged zebrafish. Dorsal and ventral slow muscles share the same processes such as myofibril assembly and muscle tissue development. On the larvae slide, 3 clusters were obtained, GO enrichment analysis suggest active muscle formation in zebrafish larvae.

## 1. Introduction

Skeletal muscle accounts for a significant proportion of vertebrate body weight, and teleost is certainly no exception. In fish, skeletal muscle can reach 35%-60% of body weight^1^. Such a high percentage of muscle mass can provide enough power for swimming, and it is also the main protein source for intensive energy activity^2^. Morphologically, the teleost trunk muscle is divided into two lateral parts by the median septum and the dorsal and ventral parts by the horizontal septum^3^. Skeletal muscle as connective tissue is composed of myotomes extending on the two sides of the body from head to tail^4^. The skeletal muscle is segmented by myosepta throughout the body into myomers which are the morphological functional units. Muscle fibers are run parallel to the body axis in the superficial region. In the deeper region, muscle fibers in a helical shape. The length of muscle fibers gets shorter as it reaches the tail. The cross-sectional area of muscle fibers is smaller in the tail and head than in the central region^4^.

Fish usually have three types of muscle fibers, which are slow-red, fast-white, and intermediate-pink muscles. Slow muscle fibers are in the narrow wedge-shaped area near to lateral line. Slow muscle fibers are also found at the base of the dorsal fins as the depressor muscle. They are a relatively small fiber diameter and consist smaller proportions of muscle tissue. They contain a large number of mitochondria^5^, store droplets, and glycogen. Their energy metabolism is exclusively aerobic^6^. They support low tail-beat frequencies and is mainly used for long and continuous swimming^7^. Fast muscle fibers consist the majority of the skeletal muscle in teleost. The diameter of the fast muscle fiber is large but decreases from head to tail. They are explosive in power and usually used for high-speed swimming, like predation and escape. The energy source is mainly the anaerobic metabolism of glycogen, but also the aerobic breakdown of lipids. The intermediate muscle fibers are located between the slow and fast muscles. As the name describes, their diameter, contraction speed, and tolerance of fatigue are between the slow and fast muscles. Some fish do not have intermediate muscle^8^, zebrafish have an intermediate muscle between slow and fast muscles.

Myogenesis involves the processes of muscle development and growth. Hyperplasia is the increase in the number of muscle fibers increases, whereas hypertrophy is the increase in the diameter of muscle fibers. According to the growth stage, myogenesis in teleosts is divided into embryonic myogenesis, stratified hyperplasia, mosaic hyperplasia, and hypertrophy. Multiple cellular events are involved in them, including activation, differentiation, proliferation, differentiation, metastasis, and fusion.

All vertebrate skeletal muscles derived from paraxial mesoderm. In the early stage of embryogenesis, the paraxial mesoderm differentiates into a somite. In the zebrafish, the paraxial mesoderm is formed by cells near to the early gastrula converging to the dorsal side. Then paraxial mesoderm is divided into segments named somites. In the early segmentation stage, there are two groups of cells in the somite, the adaxial cell and lateral presomitic cells. The adaxial cells are present as monolayers on each side of the notochord. The adaxial cells express Myogenic Regulatory Factors (MRFs), myf5 and myoD, and myosin heavy chain (MyHC) genes^9^. In the late segmentation stage, adaxial cells migrate from the notochord to myotomal surface to form the superficial slow fibers. Transcription factor Prox1 and a MyHC gene smyhc1 are expressed when adaxial cells migrate to the outer side. The lateral presomitic cells express myoD and myf5. They differentiate into medial fast muscle fibers after the adaxial cell migration. After the slow and fast muscle formation in the primary myogenesis, external cells that are separated from the epidermis appear on the lateral surface of the myotome. The external cells are called as dermomyotome. Their origin is the anterior border of the epithelial somite^10^ and they contribute to muscle growth as a source of myogenic cells^11^.

After embryonic myogenesis, hyperplasia accounts for the dominant muscle growth. New slow muscle fibers are formed at the dorsal-ventral extremities of the slow muscle layer, and new fast muscle fibers are formed at the fast muscle myotome and dorsal-ventral extremities of the myotome slow muscle layer and the side close to the slow muscle. These extreme precursor cells are present in the dermomyotome as described. They are Pax7 positive and participate in both hyperplasia and hypertrophy.

After the stratified hyperplasia, mosaic hyperplasia begins. Precursor cells are scattered among muscle cells. The precursor cells migrate deeper into the fast muscle before differentiation^12^. Small diameter new muscle fibers can be observed between existing large muscle fibers. In fast muscle, muscle fiber recruitment is ceased when fork length arrived 40% of the maximal body size ^13,14^. In slow muscle, the number of fibers continues to increase until the maximum body length^14^. Mosaic hyperplasia contributes the indeterminate growth of teleost and shows clear contrast to mammals in which hyperplastic muscle growth almost ceases at postnatal stage. In case of zebrafish, mosaic hyperplasia contributes to an approximately five-fold increase in the number of fast muscle fibers^15^.

After hyperplasia ceased, hypertrophy is the only strategy for muscle growth. the muscle fiber diameter increases. Insulin-like growth factor I (IGF-I) regulates myogenesis and muscle fiber hypertrophy. The signal regulating the hypertrophy is linked through the PI3K/AKT/mTOR pathway. IGF-I activates the PI3K/AKT/mTOR pathway, a phosphorylation cascade to increase the protein synthesis, resulting in the hypertrophy^15^. Expression of some MyHC genes is changed during the switching of mosaic hyperplasia and hypertrophy. In torafug Takifugu rubripes, a MyHC gene named MYHM2528-1 is specifically expressed in newly formed muscle fibers by hyperplasia. Transgenic fish, which expresses GFP under the regulation of MYHM2528-1 promoter, shows decreased GFP expression with the cessation of mosaic hyperplasia^16^. In zebrafish, a MyHC gene named myhz1.2 is expressed in newly formed small fibers by hyperplasia and five fast-type MyHC genes including myhz1.2, they are tandemly repeated on chromosome 5, are downregulated at the mRNA level with the cessation of mosaic hyperplasia^17^.

As described, the teleost has skeletal muscles that are morphologically different from those of mammals, such as the anatomical separation of slow and fast muscles. In previous studies, however, teleost muscle structure has been described mainly by morphological and physiological features based on the expression of a limited number of proteins, transcripts, and metabolites. In addition to structural features, teleost muscle shows unique growth phenotype. In contrast to mammalian muscle growth in which the hyperplasia process ceases after birth, the teleost exhibits continuous hyperplasia even after maturity. In zebrafish, the number of fast muscles increases rapidly at a body length of 8-10 mm, hyperplasia decreases when the body length reaches 15-17 mm, and almost stops when the body length reaches 25 mm^18^. Several transcriptional networks are involved in the continuous hyperplasia in teleost muscle^16^, detailed mechanisms are still unclear. Spatial transcriptomics is a powerful tool to obtain a comprehensive atlas of known and unknown cellular architecture of tissues. Here, we analyzed teleost muscle structure and growth characteristics from a new perspective utilizing spatial transcriptomics.

## 2. Results

### 2.1 Adult zebrafish trunk muscle clusters based on transcriptome

Tissue sections of adult fish covered 1,357 spots (supplementary 1). The spots were clustered by similarity in the whole gene expression pattern and divided into 10 clusters (Fig. 1A, C). Cluster 1 (orange) is visceral organs and cluster 2 (lime) is the skin and fins (Fig. 1A, B). Other clusters are skeletal muscle (Fig. 1A, B). Based on their location, clusters 0, 4, 5, 8, 9 and a ventral of cluster 6 are thought to be the fast muscle. Cluster 0 (red) is the largest part and distributed in 6 parts on both sides of the body. Cluster 5 (light blue) is located near the spine. Cluster 4 (spring green) and cluster 9 (cerise) are located at the dorsal and ventral regions, respectively. Spots of cluster 8 (Purple) are scattered in the fast muscle. Ventral part of cluster 6 (blue) is located at the ventral edge (Fig. 1A, B). Cluster 3 (light green) corresponds to the intermediate muscle separating the slow and fast muscles (Fig. 1A, B). However, the spots of cluster 3 are more widely extended in dorsal and ventral regions as expected. The slow muscle consists of two clusters, the dorsal part of cluster 6 (blue, dorsal slow) and cluster 7 (dark blue, lateral slow) (Fig. 1A, B).

**Figure 1.**
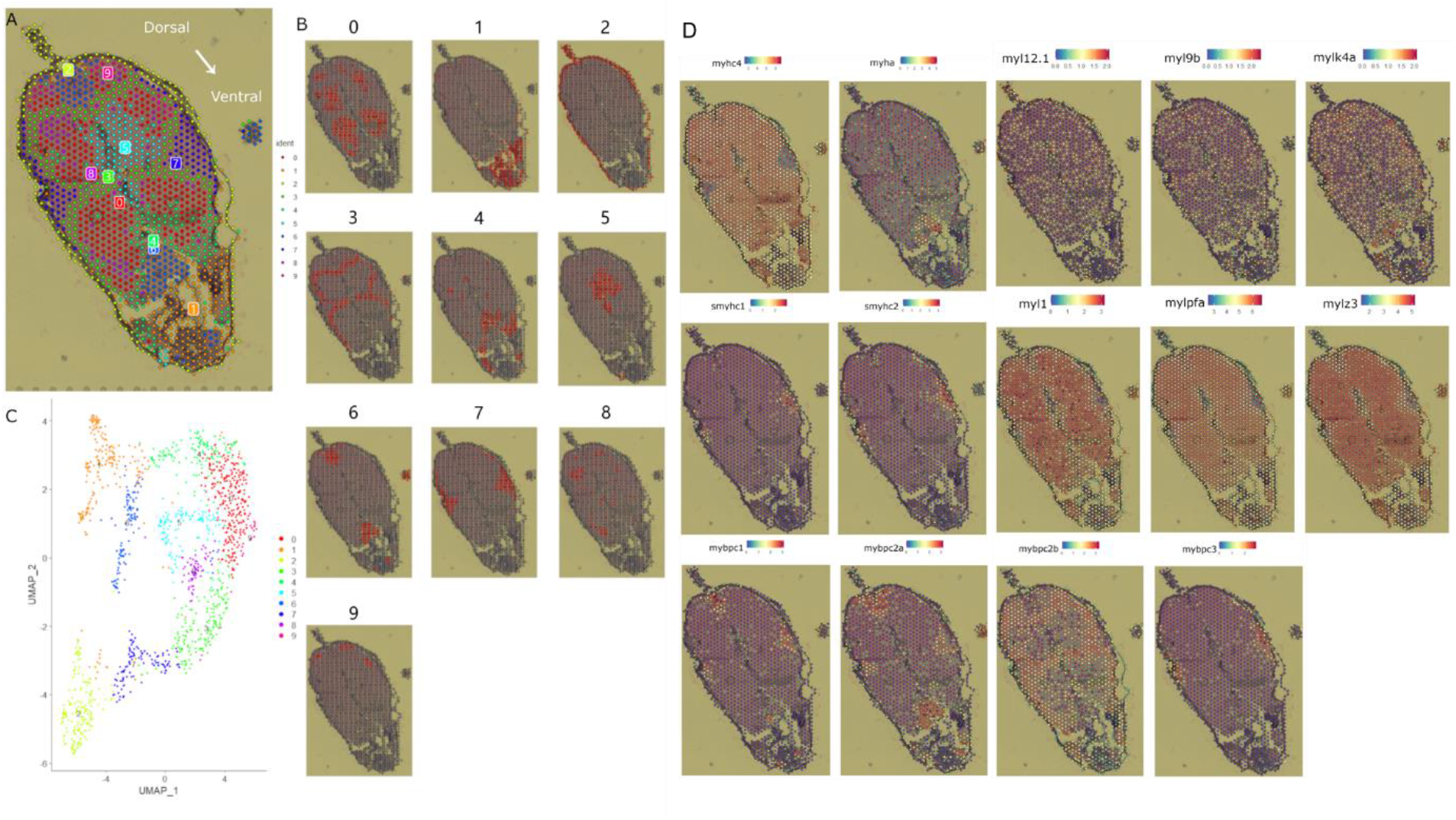
Clustering and gene expression in adult zebrafish. A, distribution of spots with cluster information. B, location of each cluster. Number indicates each cluster. C, UMAP Dimensionality Reduction (The closer the clusters are to each other, the more similar the expression is.) D, Expression of MyHC genes, myosin light chain genes, and myosin binding protein genes (Expression level in indicated with colirs. Numbers under expression color bar indicate molecular counts normalized by screansform)

### 2.2 Spatial expression of myosin heavy chain (MyHC) and other myosin related genes in adult zebrafish

Since the difference in MyHCs expression defines muscle fiber types, the spatial expression of MyHC genes was investigated and two fast-type MyHC genes, myhc4 and myha, and two slow-type MyHC genes, smyhc1 and smyhc2 were detected in our dataset (supplementary 2, Fig. 1D). myhc4 was expressed in a wide range of muscles, with the highest intensity in the fast muscle and weak expression in the lateral slow muscle. myha was mainly expressed in the ventral part of cluster 6 (Figs. 1A, B, D), indicating that the ventral part of cluster 6 is actually the fast muscle although it is clustered together with the dorsal slow muscle (dorsal part of cluster 6). Scattered expression of myha was also detected in other fast muscle regions, clusters 0, 5, and 3. The expression of smyhc1 and smyhc2 was detected in the slow muscle (Figs. 1A, B, D). smyhc1 expression was almost restricted in cluster 7 (lateral slow muscle), whereas expression of smyhc2 was detected in both lateral slow and the dorsal slow (dorsal part of cluster 6) (Figs. 1A, B, D).

Our data also showed that myosin light chain genes can be divided into two categories based on their expression. Expression of myl12.1, myl9b, and mylk4a was scattered in the whole muscle region, but mylk4a expression was high in slow muscle regions, dorsal slow and lateral slow (Fig. 1D). High expression of mylk4a was also detected in ventral part of cluster 6, despite this region is thought to be a fast muscle based on MyHC gene expression. On the other hand, myl1, mylpfa, and mylz3 were expressed predominantly in the fast muscle.

Moreover, myosin-binding protein genes showed differential expression in different muscles (Fig. 1D). mybpc2a and mybpc1 were highly expressed in cluster 6 (dorsal slow muscle and ventral fast muscle), mybpc3 was mainly expressed in cluster 7 (lateral slow muscle). mybpc2b was expressed specifically in cluster 3 (intermediate muscle) (Fig. 1D, supplementary 2).

### 2.3 GO enrichment analysis on muscle regions of adult zebrafish

GO enrichment analysis was also performed on different muscle regions of adult zebrafish (Fig. 2). In most clusters, the enriched terms are muscle structure and muscle contraction related ones. Enriched terms in cluster 5 are neural system related ones such as synaptic vesicle transport and microtuble assembly, indicating that cluster 5 contains neural cells. Cluster 3 (intermediate muscle) and cluster 7 (lateral slow) showed a similar tendency in which enriched terms are involved in the metabolic processes of nucleotide and mitochondria related pathways. Although cluster 6 expressed both slow and fast-type myosin-related genes, enriched terms of GO analysis are almost the same with the fast muscle.

**Figure 2.**
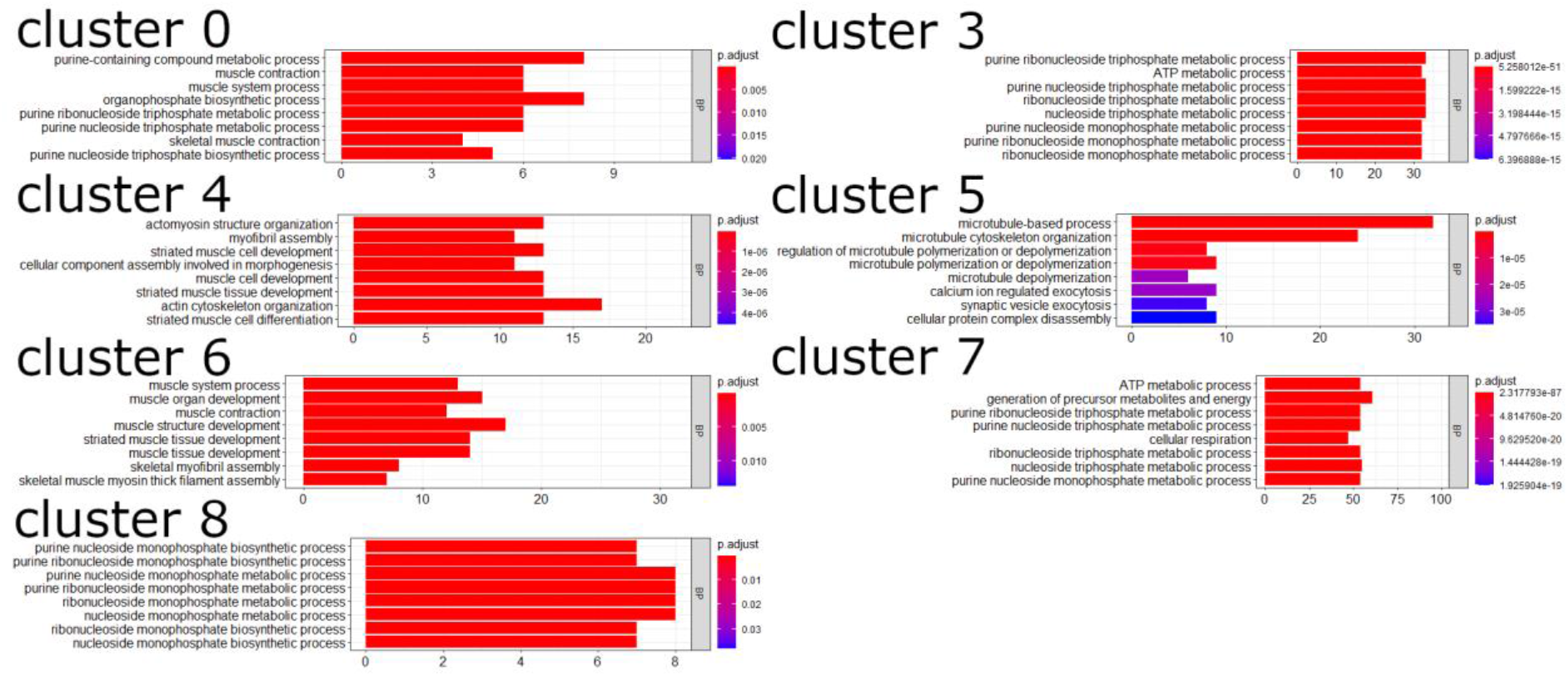
GO enrichment analysis of different muscle region in adult zebrafish.

### 2.4 Larvae zebrafish trunk muscle clusters based on transcriptome

4 sections from different individuals were subjected to Visium Spatial Transcriptome Slide. The sequencing summary is shown in Supplementary 1. Because of the small size of sections, the resolution of Visium analysis is not enough to separate the larvae muscle into different regions. Spots in each section were separated into 3 clusters, numbered 0, 1, and 2 (Fig. 3C). Cluster 0 is surface epidermal tissue (Fig. 3A, B). Marker genes of cluster 0 are epidermal tissue genes such as various keratins (supplementary 3). Spots of cluster 1 are scattered with whole body of small-sized individuals (Fig. 3A, B). Marker genes of cluster 1 contain various types of ribosomal proteins and heat shock proteins (supplementary 3), indicating active protein synthesis. Cluster 2 is the skeletal muscle. Marker genes of cluster 2 are muscle-related genes such as MyHCs, myomesin, and ryanodine receptor genes (supplementary 3).

**Figure 3.**
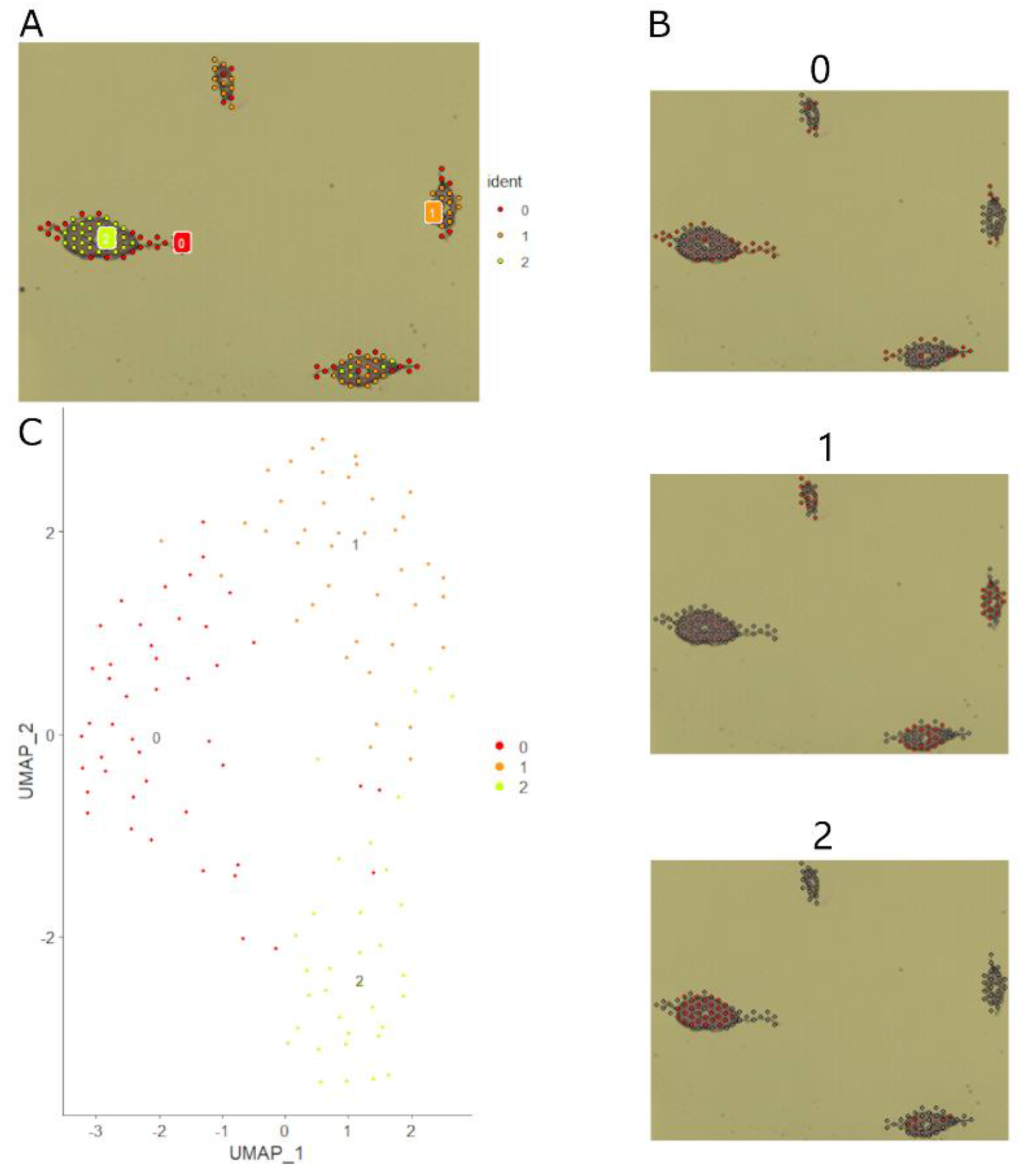
Clustering and dimensionality reduction of larvae section. A, distribution of spots with cluster information. B, location of each cluster. Number indicates each cluster. C, UMAP Dimensionality Reduction (The closer the clusters are to each other, the more similar the expression is.)

### 2.5 GO enrichment analysis on muscle regions of larvae zebrafish

Enriched terms in cluster 0 were involved in early development and tissue regeneration, such as wound healing, epiboly, fin regeneration, gastrulation, and morphogenesis of an epithelial sheet (Fig 4). Enriched terms in cluster 1 are involved in protein synthesis such as translation, peptide biosynthesis, and various metabolisms (Fig. 4). Enriched terms in cluster 2 are associated with such as muscle cell development, muscle cell differentiation, muscle structure development, myofibril assembly (Fig 4).

**Figure 4.**
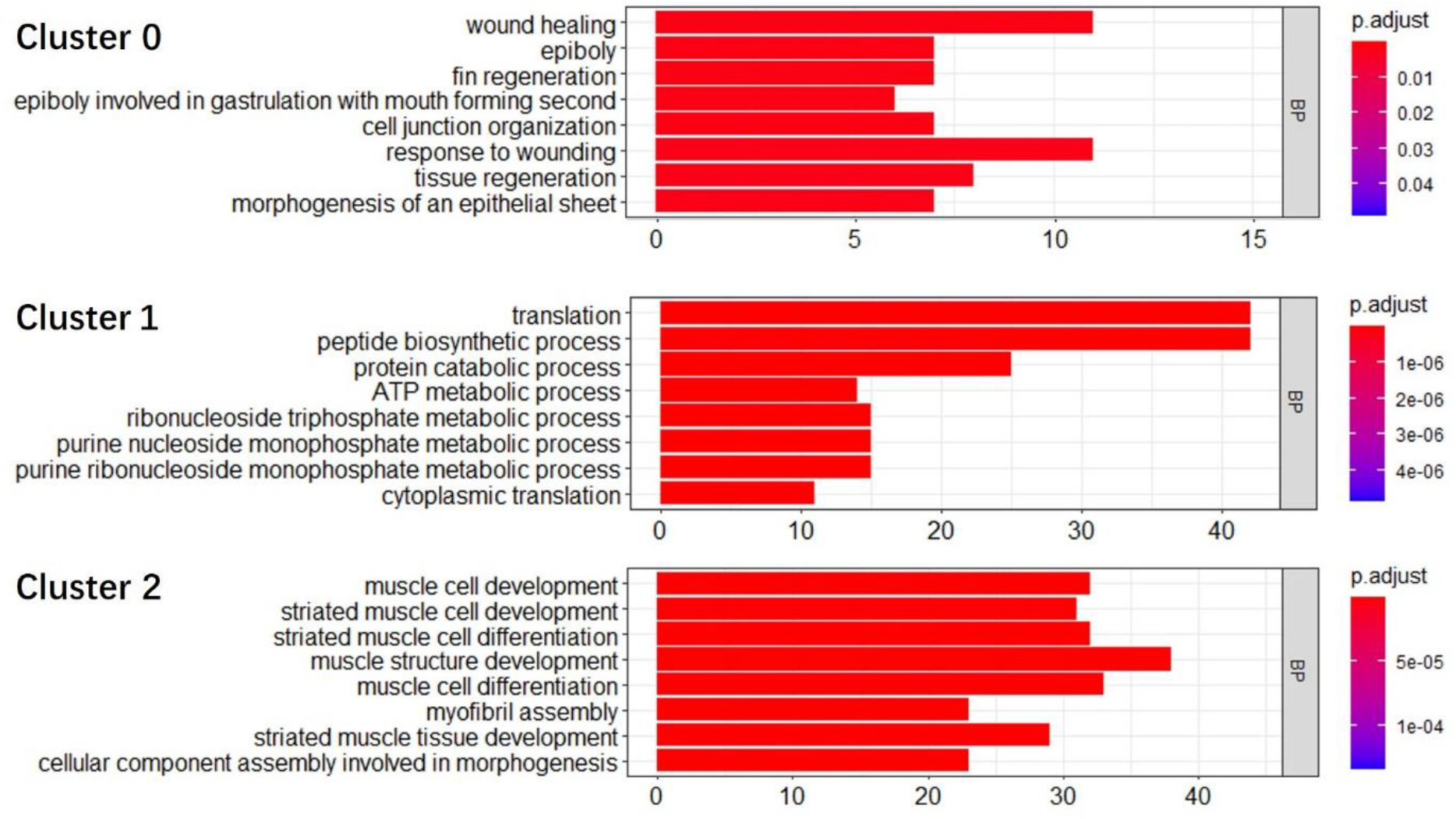
GO enrichment analysis of different muscle region in larvae zebrafish.

### 2.6 Overlapped gene markers between larvae and adult zebrafish

I examined overlap in marker genes between larvae and adult muscles. Table 1 shows the overlapped markers. Majority of overlapped markers are genes detected in the fast muscle of the adult, indicating that majority of larvae cluster 2 is the fast muscle. Marker genes of adult cluster 3 (intermediate muscle) and cluster 6 (dorsal slow and ventral fast muscle) were also detected in larvae cluster 2. As for adult lateral slow muscle (cluster 7), only acta1b was detected in the larvae muscle. On the other hand, marker genes specific to larvae muscle are listed in Table 2. MyHC gene named myhz1.3 was detected as a specific marker of larvae muscle.

**Table 1.**
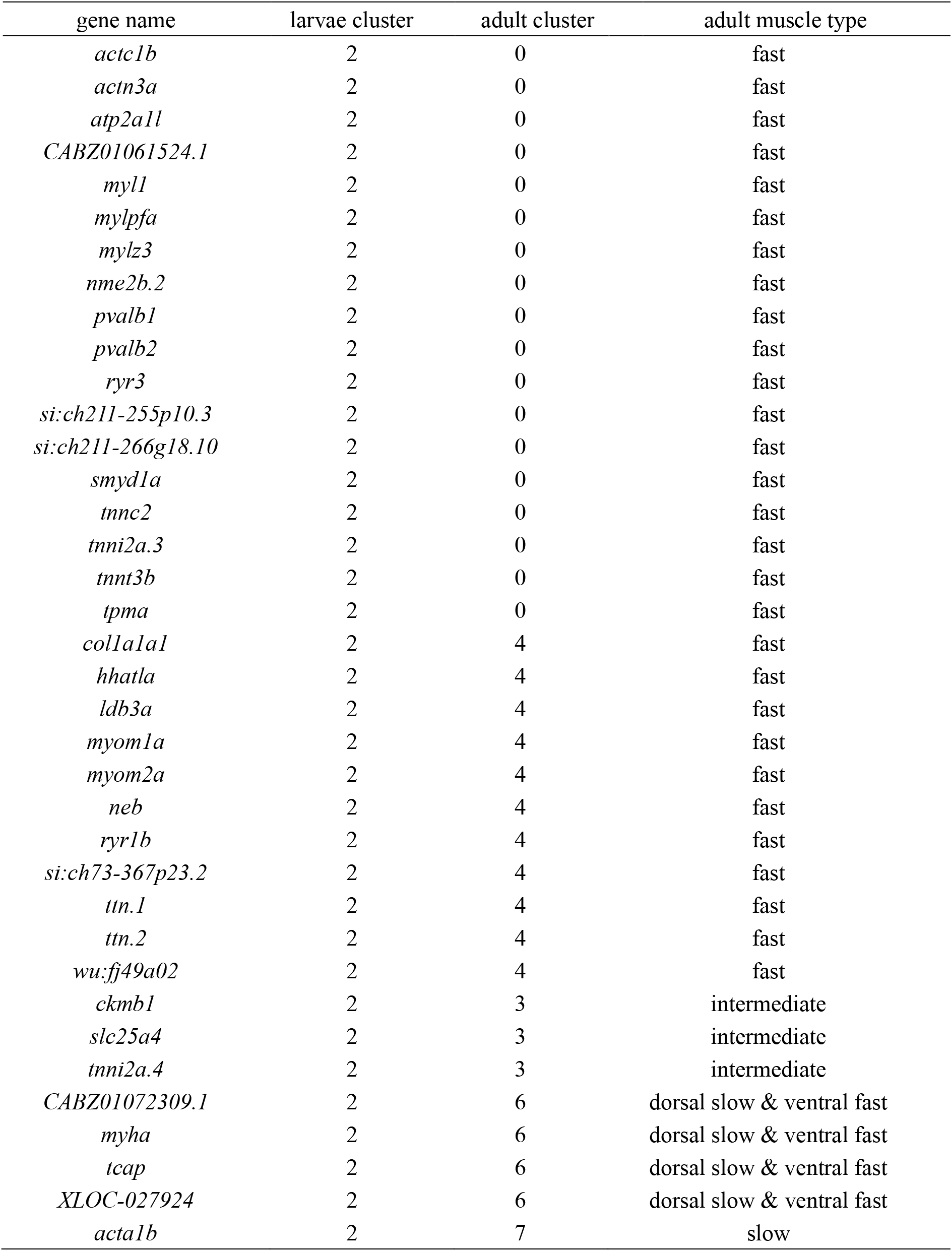
Gene markers shared by larvae and adult fish muscles

**Table 2.**
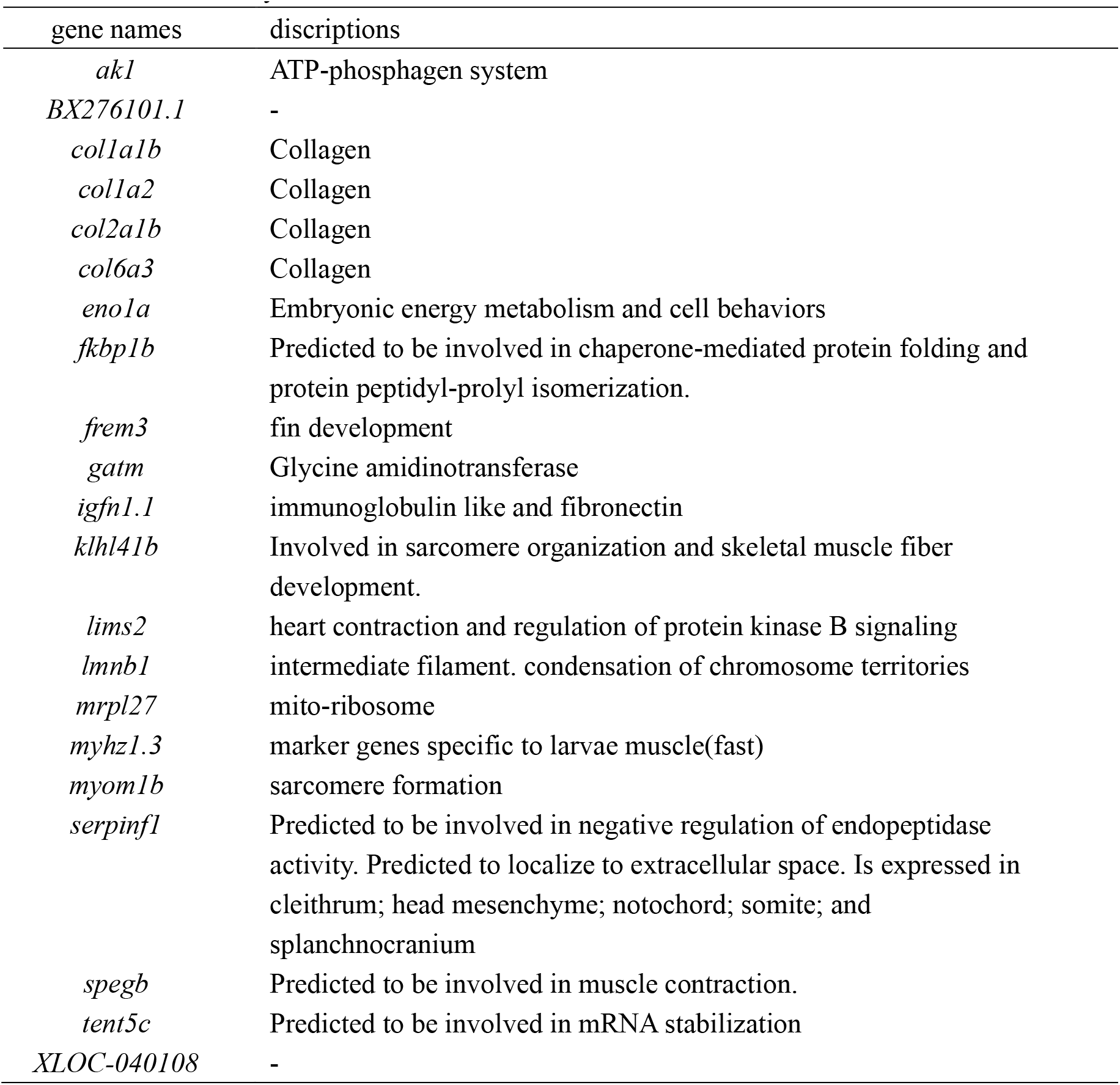
Gene marker only exist in larvae muscle

## 3. Discussion

### 3.1 Adult zebrafish trunk muscle structure

Skeletal muscles are classified into fast, slow muscles based on their contractile characteristics, which are mainly determined by the expression of MyHCs. In this study, 4 MyHC genes were detected in our spatial transciptomic dataset. smyhc1 and smyhc2 are slow-type MyHC genes. During teleost myogenesis, adaxial cells migrate from center to the surface to form slow muscle. These adaxial cells express smyhc1. Fast-type MyHC gene, myhc4 is also detected in the adaxial cell at the 7-somite stage, myhc4 is expressed only in fast muscle fibers after adaxial cell migration^19^. myha (fmyhc2.3) is also expressed in fast muscle fibers^20^. Myosin light chain genes and myosin binding protein genes also show muscle fiber-type specific expression. myl1 is a fast specific marker that is expressed earlier than others^21^. mylpfa and mylz3 are only expressed in fast muscle^22^. mybpc1 is expressed in the slow muscle, whereas mybpc2 is expressed in fast muscle. mybpc3 is expressed in cardiac muscle^23,24^.

Figure 5 shows schematics of the standard muscle structure of teleost (Fig. 5A) and clusters detected by spatial transcriptomics in this study (Fig. 5B). Cluster 0 consists the majority of the fast muscle and showed representative fast muscle-type gene expression such as fast-type MyHC genes, myosin light chain genes, and myosin-binding protein genes (Fig. 1D). Cluster 5 also shares a similar gene expression pattern with cluster 0 in MyHC and myosin-related genes, but its location was restricted near the spine. Enrichment analysis showed that muscle contraction-related pathways were upregulated in cluster 0, whereas cell movement, cytoskeleton, microtuble related pathways were upregulated in cluster 5, suggesting relation with the neural system.

**Figure 5.**
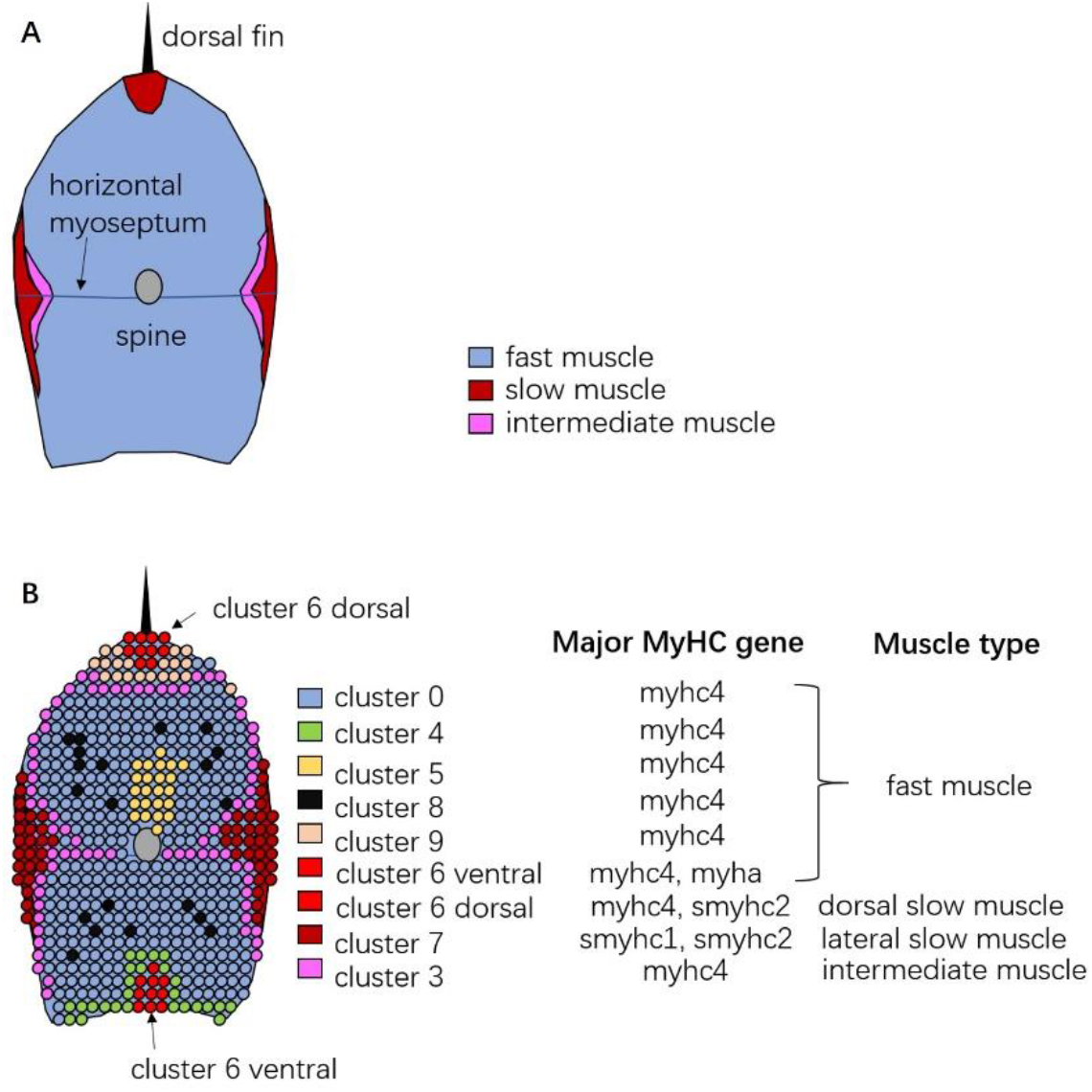
Schematics of standard composition of teleost trunk skeletal muscle (A) and clusters detected by spatial transcriptomics (B)

As for the slow muscle, co-expression of two slow-type MyHC genes, smyhc1 and smyhc2, was observed in cluster 7 (Fig. 1D). Fast-type MyHC gene expression was clearly downregulated in cluster 7. Combining with the location, it is clear that cluster 7 is the lateral slow muscle. Enrichment terms also showed upregulation of mitochondrial respiration related pathways, a representative physiological character of the slow muscle.

Classification of cluster 6 is more complex. Cluster 6 is divided into two parts, the ventral part and dorsal part (cluster 6 ventral and cluster 6 dorsal in Fig. 1D). MyHC gene expression was clearly different between the two parts, the ventral part expressed fast-type myhc4 and myha but not slow-type MyHC genes, whereas the dorsal part expresses slow-type smyhc1 and smyhc2 and fast-type myhc4. The dorsal part of cluster 6 had similarities to the lateral slow muscle (cluster 7) in terms of expressions of myosin light chain and myosin binding protein genes (Fig. 1D). Combining with the location, the the dorsal part of cluster 6 is dorsal slow muscle, and ventral part of cluster 6 is the fast muscle. However, it is unclear whether fast and slow fibers are mixed in these regions or not.

Cluster 3 is located between slow and fast muscles, indicating that cluster 3 is the intermediate muscle. The expression pattern of myosin-associated genes in cluster 3 was almost the same with the fast muscle, except for mybpc2b, which was expressed specifically in cluster 3 (Fig. 1D, supplementary 2). This result suggests that mybpc2b can be a useful marker to identify intermediate muscle. On the other hand, enrichment analysis showed that the intermediate muscles are similar to the lateral slow muscle (cluster 7), and various energy metabolic processes and mitochondrial respiration processes are upregulated in the intermediate muscles. Intermediate muscle, as its name suggest, is intermediate between slow and fast muscle even at transcriptome levels.

Muscle development terms were enriched in two fast muscle-related clusters, cluster 4 and cluster 6 (Fig. 2). Cluster 4 belongs to the fast muscle due to the expression of fast type MyHC. Cluster 6 contains dorsal slow and ventral fast muscles. As described, teleost muscle growth continues in the slow muscle but ceases in the fast muscle. This is unexpected results because in previous studies, the growth of fast muscles in zebrafish stagnated after it reaches about 25 mm total length. Size of zebrafish used in this study was about 30 mm. However, our results suggest that the dorsal and ventral edge including fast muscle continue the muscle growth even at the aged stage.

### 3.2. Growth Characteristics of Zebrafish Muscle and Spatial Transcriptomics

It is clear that myogenic activity is high in larvae due to enriched terms of muscle differentiation and muscle growth in larvae muscle (Fig. 4) and myod1 expression in the whole myotomal region of larvae (Fig. 6). On the other hand, such upregulation was detected only in a limited region (or scattered spots) of adult muscle (Fig. 6). Taken together with upregulation of Kelch family genes, which is involved in the proliferation and differentiation of myogenic cells, myogenic acidity in larvae is high in comparison with adult. myhz1.2 is considered a marker of newly formed muscle fibers by hyperplasia^17^. Expression of myhz1.2 was detected in whole myotomal region of larvae, indicating active hyperplastic growth in larvae. Expression of myhz1.2 was also detected in adult but their number is very few and the distribution is scattered (Fig. 6).

**Figure 6.**
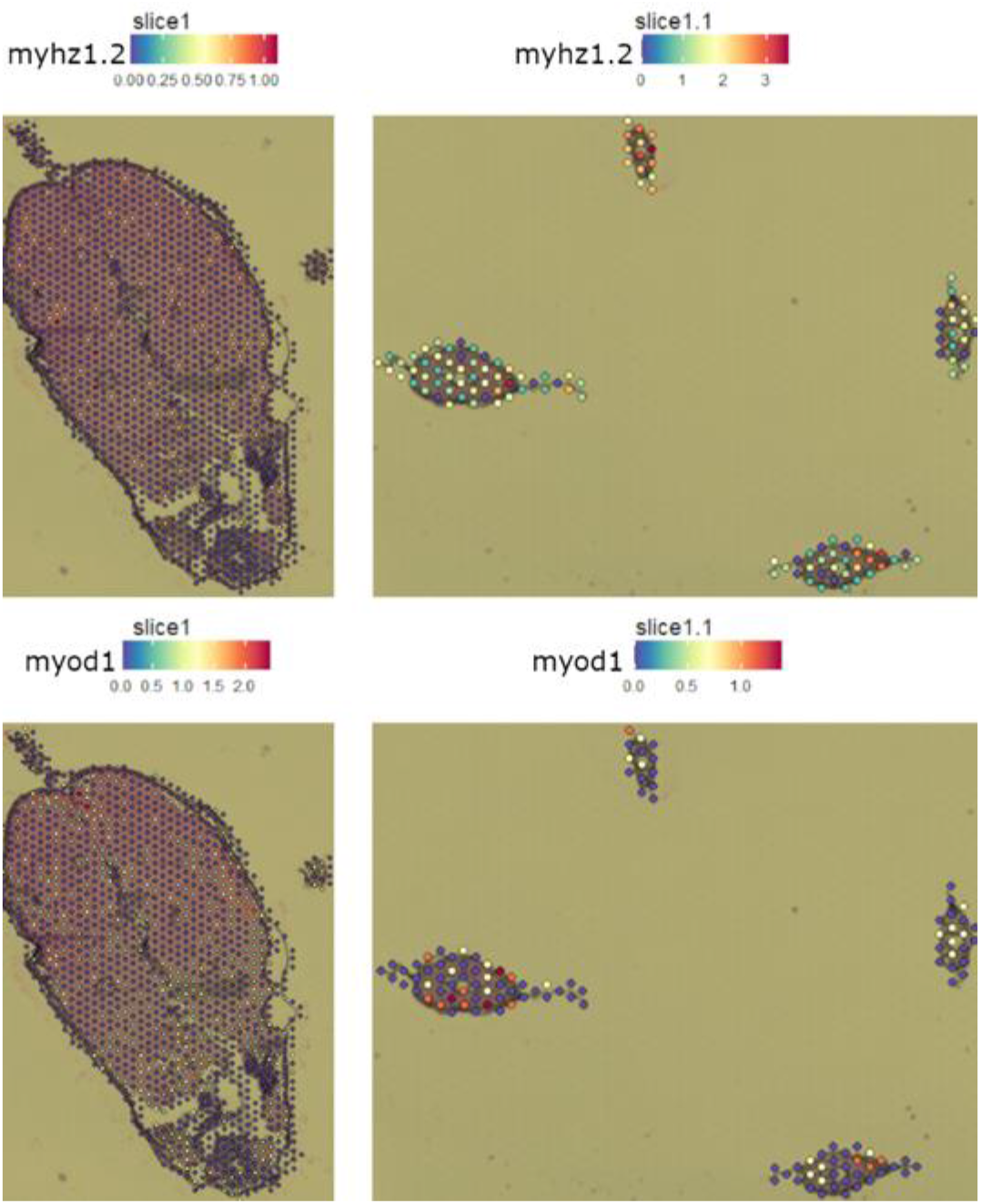
myhz1.2 (upper panel) and myod1 (lower panel) expression in adult and larvae zebrafish. Expression level is indicated with colors. Numbers under expression color bar indicate molecular counts normalized by sctransform

Interesting feature identified in this study is the myogenic activity of a part of fast muscle in the adjacent narrow areas of the ventral adult musculature (cluster 4 and cluster 6 ventral, see Fig. 5). As described, myogenesis is downregulated in fast muscle of matured adult fish. In this region, however, despite clear expression of fast-type MyHC genes, myhc4, myog was also upregulated and its expression is higher than that in the slow muscle region. myog is thought to regulate the late step of the muscle differentiation. Actually, the enriched biological processes in this region contained muscle cell development, muscle cell differentiation (Fig. 2), revealing that the muscle growth process is upregulated in ventral region of the fast muscle. The possibility that the ventral part of the fast muscle is a center of myogenesis at adult stage is interesting. myha, a major fast MyHC gene expressed in larvae muscle (supplementary 3, table 1), was also detected in the ventral fast (Fig. 1D), indicating similarities between ventral fast and larvae muscle which shows active myogenesis. Additional examination is required to examine this possibility. Stem cell number and activity are also important to examine ability of myogenesis. Most transcription factors especially muscle stem cell markers such as pax3 and pax7, however, were not detected in this study due to their low expression levels.

## 4. Materials and methods

### 4.1 Animal care

Zebrafish RW strain obtained from Riken was cultured in a 28°C constant temperature filtered recirculating water system that provided light from 9:30 am to 9:30 pm. Powdered feed was fed daily in the morning and paramecia to juveniles and brine shrimp to adults in the afternoon on weekdays. Feeding once a day on weekends and holidays. Clean filters daily and change them regularly. Eggs were obtained by breeding with adults over 6 months of age. Three pairs of parents were placed in the breeding tank the night before, and the males and females were separated by a partition. The next day, when the light was provided, the partitions were removed and the eggs were collected and raised in methyl blue solution in a 28°C incubator until they hatched, and the larvae were transferred to the system after 5 days. In this study, adult zebrafish were about 8 months old and 4 larvae were about 1 month old(supplementary 4).

### 4.2 Spatial Transcriptomics

We employed the Visium Spatial Gene Expression solution (10X Genomics). On a Visium Spatial Gene Expression slide, there are 4 capture areas with 5,000 barcode spots in each capture area. Primers were present on each barcode spot and comprised of Illumina TruSeq Read1, 16 nt Spatial Barcode, 12 nt unique molecular identifiers, and 30 nt poly(dT) sequence (supplementary 5). The obtained cDNA library contains information on the location of the spots and is capable of overlaying gene expression with bright-field micrographs after processing. In this study, two capture zones were used, one for adults and one for 4 larvae zebrafish.

Zebrafish were anesthetized in ice water, and an appropriate length of the trunk was intercepted forward from the anal. After removing blood using a wiper, the tissues were covered with an optimal cutting temperature compound (OCT compound, SAKURA). Tissues were transferred to the mold (SAKURA) and covered by OCT compound. The sample was immersed in an isopentane-liquid nitrogen bath until the surface becomes completely white and opaque. Place Visium slide, Tissue Optimization slide, and tissue blocks into the cryostat (SAKURA) chamber for 30 min for temperature equilibrium. Section thickness was set to 10 μm. The tissue blocks were then sliced and mounted within the fiducial frame on the slides. Transfer slides on dry ice to -80°C for storage.

Methanol was chilled to -20. Tris-Acetic Acid Buffer (11 g Tris base in 200 ml nuclease-free water, pH 6.0, filtered through 0.2 μm nylon membrane filter) was prepared before the experiment. Thermocycler Adaptor was placed on the thermal cycler set at 37°C for 5 mins. Slide incubated for 1 min. The slide was fixed in pre-cooled methanol at -20°C for 30 mins.

Milli-Q water and Eosin mix (100 μl/slide Eosin Y solution and 900 μl/slide Tris-Acetic Acid Buffer) were prepared. Remove the slide from methanol. Add 500 μl isopropanol for 1 min. Remove the reagent and air dry the slide in less than 10 min. Add 1 ml Hematoxylin for 7 min. Rinse in the Milli-Q water. Added 1 ml Bluing buffer for 2 min. The reagent was discarded, and the slide was rinsed in the Milli-Q water. Added 1 ml Eosin Mix, incubated for 1 min. Discard the reagent. The slide was washed with Milli-Q water. Incubated at 37°C for 5 mins. Taking microscope (KEYENCE) photos in a bright field.

Equilibrate the reagents. Setting the slide in the slide cassette, 70 μl permeabilization enzyme was added to each well, placing the cassette on the Thermocycler adaptor at 37°C, the permeabilization time was determined in a preliminary experiment by using Visium Spatial Tissue Optimization slide, here, the optimization time was 6 min (supplementary 6). Remove Permeabilization enzyme from each well. Add 100 μl 0.1X SSC to the wells.

For reverse transcription, prepare RT Master Mix on ice. Remove 0.1X SSC, add 75 μl RT Master Mix to each well. Put the sealed cassette on the thermocycler and run the program (lid temperature at 53, 53 45 min, 4°C hold).

For second strand synthesis, remove RT Master Mix from the walls, add 75 μl 0.08 KOH, incubated for 5 min at room temperature. Remove KOH, add 100 μl EB to each well. Prepare Second Strand Mix, Add 75μl to each well. Put the sealed cassette on the thermocycler and run the program (lid temperature at 65°C, 65°C 15 min, 4°C hold). Remove the reagents and add 100 μl EB to each well. Remove Buffer EB and add 35 μl 0.08M KOH, incubate for 10 min at room temperature. Add 5 μl Tris (1 M, pH 7.0) to 4 tubes in an 8-tube strip and Transfer 35 μl sample from each well to a corresponding tube containing Tris in the 8-tube strip. Cycle number was determined by qPCR before performing cDNA amplification. Mix 1 µl of sample and 9 μl qPCR Mix, run the program (98°C 3 min,98°C 5 s, 63°C 30 s, read signal, to step2, 25 cycles), The threshold for determining the Cq Value should be set along with the exponential phase of the amplification plot, at ∼25% of the peak fluorescence value. According to the plot, adult zebrafish were set to 19 cycles, and juvenile fish, due to the small size of the sample, were increased by two cycles to 21 cycles compared to adult fish.

Add 65 μl cDNA amplification Mix to the remaining sample, run the thermal cycler with the program (lid temperature 105°C, 98°C 3 min, 98°C 15s, 63°C 20s, 72°C 1 min 19 cycles for adult sample 21 cycles for larvae sample, 4°C hold). The sample was stored at 4°C.

For clean-up, Add 60 μl 0.6X SPRIselect reagent, incubate for 5 min, place on the magnet, and remove the supernatant solution clear. Add 200 μl 80% ethanol. Remove ethanol. Wash with ethanol again. Air dry for 2 min. add 40.5 μl Buffer EB, incubate 2 min. Place the sample on the magnet, transfer 40 μl sample to a new tube strip. Run 1 μl of sample on Agilent High Sensitivity tape for Quantification.

Pre-cool the thermal cycler to 4°C. Prepare Fragmentation Mix on ice. Transfer 10 μl purified cDNA sample to tube strips on ice. Add 25 μl Buffer EB and 15 μl Fragmentation mix to each sample. Transfer the tubes into a precooled thermal cycler and run program (lid 65°C,32°C 5 min, 65°C 30 min, 4°C hold).

Add vortexed 0.6X SPRIselect 30 μl to each sample, incubated for 5 mins. Place on the magnet, transfer 75 μl supernatant to the new tube strip. Add vortexed 0.8X SPRIselect 10 μl to each sample, incubated for 5 mins. Place on the magnet, remove 80 μl supernatant. Add 125 μl 80% ethanol, remove ethanol after 30 s. Wash it twice. Add 50.5 μl Buffer EB to each sample and incubate 2 mins. Place on the magnet then transfers the 50 μl sample to the new tube strip after solution clear.

Prepare Adaptor Ligation Mix, Mix 50 μl Adaptor Ligation Mix, and 50 μl sample. Run the thermal cycler with the program(lid temperature 30°C, 20°C 15 min, 4°C hold). Add vortexed 0.8X SPRIselect 80 μl to each sample, incubated for 5 mins. Place on the magnet, remove the supernatant. Add 200 μl 80% ethanol, remove ethanol after 30 s. Wash it twice. Add 30.5 μl Buffer EB to each sample and incubate 2 mins. Place on the magnet then transfers the 30 μl sample to the new tube strip after solution clear.

Add 50 μl Amp Mix and 20 μl Dual Index TT Set A to sample. Run thermal cycler with the program(lid temperature 105°C, 98°C 45 s, 98°C 20s, 54°C 30s, 72°C 20 s 14 cycles, 72°C 1min, 4°C hold). Add vortexed 0.6X SPRIselect 60 μl to each sample, incubated for 5 mins. Place on the magnet, transfer 150 μl supernatant to the new tube strip. Add vortexed 0.8X SPRIselect 20 μl to each sample, incubated for 5 mins. Place on the magnet, remove 165 μl supernatant. Add 200 μl 80% ethanol, remove ethanol after 30 s. Wash it twice. Add 35.5 μl Buffer EB to each sample and incubate 2 mins. Place on the magnet then transfers 35 μl sample to a new tube strip after solution clear(supplementary 7).

The sample was sequenced with HiSeqXTen. The sequencing depth for the adult sample was 55 million and the depth for larvae was 3 million by considering the coverage of the tissue section(supplementary 8).

### 4.2 Bioinformation tools

We used Space Ranger (10X Genomics) and Seurat (Satija Lab).

This pipeline is provided by 10X genomics. Annotation file V4.2 (Lawson et al., 2020) was used to avoid a 3’ bias of Visium. Reference data files could be prepared with spaceranger mkref. Spaceranger mkfastq merge Illumina’s bcl2fastq files into FASTQ files and subsequently Spaceranger count to combine the resulting FASTQ with bright field microscopy images to pair the sequences, barcode, UMI count, etc. The output spot feature matrix, clustering, and gene expression analysis were performed. (10X Genomics)

Seurat is an R package for analyzing single-cell RNA sequencing data. The spatial transcriptome dataset is like that of single-cell sequencing, and tools focusing on spatial visualization have been added to this package, In addition to the above, the following package to also be used: ggplot2, patchwork, dplyr, RColorBrewer, clusterProfiler, etc. Therefore, functions such as normalization, dimensionality reduction, clustering, and GO enrichment analysis can be accomplished with them. ^25,26,27,28,29,30^

## Supporting information

Sequence information

Marker genes of each cluster in adult zebrafish muscle

Marker genes of each cluster in larvae zebrafish muscle

experimental materials

Visium method

Permeabilization optimization and Quality check of cDNA libraries

## Acknowledgments

This research was partially funded by JSPS KAKENHI Grant Number 20H00427.

The authors have no conflicts to disclose.

