## Supplementary material for "Zebrafish *Danio rerio* trunk muscle structure and growth from a spatial transcriptomics perspective": Sequence information

supplementary 1 Sequence result summary

| Terms | Adult | Larvae |
|---|---|---|
| Spots under tissue | 1,357 | 120 |
| Mean reads per spot | 134,320 | 194,582 |
| Median gene per spot | 1,243 | 2,122 |
| Total genes detected | 22,388 | 16,973 |
| Median UMI Counts per spot | 6,220 | 8,092 |
