## Supplementary material for "Zebrafish *Danio rerio* trunk muscle structure and growth from a spatial transcriptomics perspective": Marker genes of each cluster in adult zebrafish muscle

supplementary 2 Marker genes of each cluster in adult zebrafish muscle ($p<0.05$)

| cluster | muscle type | marker gene | avg_logFC | p_val_adj (<0.05) |
|---|---|---|---|---|
| 0 | fast | *mylpfa* | 0.493 | 2.03E-83 |
| 0 | fast | *pvalb2* | 0.929 | 2.53E-82 |
| 0 | fast | *pvalb3* | 0.554 | 1.58E-80 |
| 0 | fast | *actc1b* | 0.425 | 3.55E-78 |
| 0 | fast | *ckma* | 0.469 | 3.77E-76 |
| 0 | fast | *mylz3* | 0.511 | 3.42E-74 |
| 0 | fast | *tnnt3b* | 0.434 | 1.26E-67 |
| 0 | fast | *ckmb* | 0.394 | 4.77E-64 |
| 0 | fast | *pvalb1* | 0.574 | 3.42E-62 |
| 0 | fast | *mybphb* | 0.674 | 5.54E-62 |
| 0 | fast | *tpma* | 0.446 | 5.85E-62 |
| 0 | fast | *si:dkey-151g10.6* | 0.415 | 2.06E-56 |
| 0 | fast | *nme2b.2* | 0.344 | 4.24E-56 |
| 0 | fast | *myl1* | 0.523 | 9.36E-53 |
| 0 | fast | *si:dkey-51e6.1* | 0.447 | 4.59E-43 |
| 0 | fast | *rpl38* | 0.373 | 1.44E-37 |
| 0 | fast | *ampd1* | 0.434 | 2.28E-37 |
| 0 | fast | *tnni2a.3* | 0.317 | 2.56E-37 |
| 0 | fast | *actn3a* | 0.408 | 8.67E-33 |
| 0 | fast | *atp2a1l* | 0.345 | 1.09E-32 |
| 0 | fast | *CABZ01078594.1* | 0.483 | 4.89E-30 |
| 0 | fast | *pvalb4* | 0.28 | 2.03E-27 |
| 0 | fast | *tnnc2* | 0.298 | 2.97E-27 |
| 0 | fast | *myoz1a* | 0.383 | 1.21E-26 |
| 0 | fast | *rps28* | 0.271 | 8.79E-26 |
| 0 | fast | *aldoab* | 0.304 | 1.03E-22 |
| 0 | fast | *si:dkey-211f22.5* | 0.415 | 5.09E-22 |
| 0 | fast | *smyd1a* | 0.301 | 2.57E-21 |
| 0 | fast | *rpl27a* | 0.31 | 4.97E-21 |
| 0 | fast | *pkmb* | 0.299 | 3.66E-20 |
| 0 | fast | *rpl9* | 0.332 | 5.68E-20 |
| 0 | fast | *atp5f1e* | 0.289 | 1.57E-19 |
| 0 | fast | *lyrm2* | 0.304 | 6.41E-17 |
| 0 | fast | *aldoaa* | 0.253 | 7.97E-17 |
| 0 | fast | *uqcrb* | 0.317 | 1.02E-16 |
| 0 | fast | *prr33* | 0.335 | 1.31E-16 |
| 0 | fast | *gamt* | 0.331 | 1.93E-16 |
| 0 | fast | *tmod4* | 0.293 | 4.34E-16 |
| 0 | fast | *CABZ01061524.1* | 0.292 | 9.90E-15 |
| 0 | fast | *myoz1b* | 0.254 | 1.76E-14 |

| cluster | muscle type | marker gene | avg_logFC | p_val_adj (<0.05) |
|---|---|---|---|---|
| 0 | fast | *CABZ01070747.1* | 0.275 | 1.80E-13 |
| 0 | fast | *si:ch211-255p10.3* | 0.256 | 3.55E-13 |
| 0 | fast | *casq1a* | 0.285 | 3.82E-13 |
| 0 | fast | *si:ch211-266g18.10* | 0.33 | 4.70E-13 |
| 0 | fast | *smdt1a* | 0.258 | 1.61E-12 |
| 0 | fast | *gyg1b* | 0.252 | 2.59E-12 |
| 0 | fast | *ryr3* | 0.255 | 8.81E-12 |
| 0 | fast | *txlnbb* | 0.316 | 4.61E-11 |
| 0 | fast | *rtn4b* | 0.25 | 6.89E-11 |
| 3 | intermediate | *eno3* | 0.556 | 3.13E-53 |
| 3 | intermediate | *ND3* | 0.712 | 4.28E-52 |
| 3 | intermediate | *ATP6* | 0.639 | 4.28E-52 |
| 3 | intermediate | *ND4* | 0.586 | 7.16E-43 |
| 3 | intermediate | *ND2* | 0.591 | 8.42E-42 |
| 3 | intermediate | *aldoaa1* | 0.477 | 3.66E-41 |
| 3 | intermediate | *mt-cyb* | 0.433 | 5.00E-41 |
| 3 | intermediate | *mt-co1* | 0.565 | 1.18E-40 |
| 3 | intermediate | *slc25a5* | 0.518 | 4.19E-40 |
| 3 | intermediate | *mt-co2* | 0.48 | 1.92E-39 |
| 3 | intermediate | *mt-nd1* | 0.572 | 4.22E-39 |
| 3 | intermediate | *gapdh* | 0.304 | 1.18E-38 |
| 3 | intermediate | *atp5f1e1* | 0.491 | 2.96E-36 |
| 3 | intermediate | *NC-002333.17* | 0.55 | 7.63E-36 |
| 3 | intermediate | *pvalb41* | 0.387 | 9.92E-35 |
| 3 | intermediate | *ybx1* | 0.39 | 1.48E-34 |
| 3 | intermediate | *mybpc2b* | 0.626 | 1.15E-33 |
| 3 | intermediate | *COX3* | 0.396 | 7.25E-33 |
| 3 | intermediate | *tpi1b* | 0.472 | 8.99E-32 |
| 3 | intermediate | *atp5mc3b* | 0.364 | 8.99E-30 |
| 3 | intermediate | *atp5md* | 0.447 | 2.44E-27 |
| 3 | intermediate | *mt-nd5* | 0.462 | 5.94E-27 |
| 3 | intermediate | *ndufa3* | 0.48 | 3.41E-25 |
| 3 | intermediate | *nme2b.21* | 0.296 | 5.36E-25 |
| 3 | intermediate | *pgam2* | 0.434 | 7.14E-25 |
| 3 | intermediate | *atp5if1a* | 0.519 | 3.86E-24 |
| 3 | intermediate | *slc25a4* | 0.345 | 4.08E-23 |
| 3 | intermediate | *NC-002333.14* | 0.424 | 8.47E-23 |
| 3 | intermediate | *tnni2a.4* | 0.503 | 3.22E-22 |
| 3 | intermediate | *idh2* | 0.315 | 6.31E-22 |
| 3 | intermediate | *mdh2* | 0.343 | 7.04E-22 |
| 3 | intermediate | *zgc:193541* | 0.378 | 4.87E-20 |
| 3 | intermediate | *trnS2* | 0.361 | 5.10E-19 |

| cluster | muscle type | marker gene | avg_logFC | p_val_adj (<0.05) |
|---|---|---|---|---|
| 3 | intermediate | *atp5l* | 0.342 | 6.49E-19 |
| 3 | intermediate | *cox7c* | 0.337 | 4.19E-18 |
| 3 | intermediate | *atp5f1c* | 0.358 | 1.14E-17 |
| 3 | intermediate | *cox6a2* | 0.303 | 1.48E-17 |
| 3 | intermediate | *ndufb7* | 0.354 | 2.13E-16 |
| 3 | intermediate | *atp5pd* | 0.345 | 2.83E-16 |
| 3 | intermediate | *NC-002333.41* | 0.329 | 7.66E-16 |
| 3 | intermediate | *ndufa1* | 0.372 | 1.85E-15 |
| 3 | intermediate | *atp5po* | 0.313 | 5.40E-15 |
| 3 | intermediate | *ckmb1* | 0.274 | 7.86E-15 |
| 3 | intermediate | *tnni2b.2* | 0.359 | 1.01E-14 |
| 3 | intermediate | *ndufa7* | 0.356 | 1.03E-14 |
| 3 | intermediate | *cox6c* | 0.301 | 3.53E-14 |
| 3 | intermediate | *aldoab1* | 0.303 | 4.76E-14 |
| 3 | intermediate | *uqcrq* | 0.362 | 6.99E-14 |
| 3 | intermediate | *uqcr10* | 0.381 | 8.22E-14 |
| 3 | intermediate | *ndufa2* | 0.375 | 1.31E-13 |
| 3 | intermediate | *CABZ01102240.1* | 0.328 | 2.33E-13 |
| 3 | intermediate | *uqcrb1* | 0.411 | 6.45E-13 |
| 3 | intermediate | *atp5f1d* | 0.326 | 1.86E-12 |
| 3 | intermediate | *cox5b2* | 0.267 | 1.86E-12 |
| 3 | intermediate | *cox7a1* | 0.323 | 1.87E-12 |
| 3 | intermediate | *atp5pb* | 0.311 | 2.91E-12 |
| 3 | intermediate | *NDUFB1* | 0.302 | 3.97E-12 |
| 3 | intermediate | *hbaa1* | 0.403 | 4.40E-12 |
| 3 | intermediate | *ckma1* | 0.291 | 8.09E-12 |
| 3 | intermediate | *ndufb3* | 0.377 | 9.76E-12 |
| 3 | intermediate | *atp5pf* | 0.318 | 1.51E-11 |
| 3 | intermediate | *ckmt2a* | 0.318 | 2.50E-11 |
| 3 | intermediate | *ndufb4* | 0.34 | 1.15E-10 |
| 3 | intermediate | *cox6b2* | 0.291 | 3.30E-10 |
| 3 | intermediate | *vdac2* | 0.277 | 4.09E-10 |
| 3 | intermediate | *ndufa4l* | 0.275 | 7.66E-10 |
| 3 | intermediate | *ND4L* | 0.274 | 7.74E-10 |
| 3 | intermediate | *slc25a11* | 0.292 | 1.82E-09 |
| 3 | intermediate | *atp5meb* | 0.296 | 9.85E-09 |
| 3 | intermediate | *atp5mc1* | 0.28 | 1.11E-08 |
| 3 | intermediate | *ckmt2b* | 0.253 | 1.16E-08 |
| 3 | intermediate | *mpc1* | 0.273 | 1.18E-08 |
| 3 | intermediate | *ndufs4* | 0.255 | 3.18E-08 |
| 3 | intermediate | *fh* | 0.283 | 3.45E-08 |
| 3 | intermediate | *pgm1* | 0.292 | 3.87E-08 |

| cluster | muscle type | marker gene | avg_logFC | p_val_adj (<0.05) |
|---|---|---|---|---|
| 3 | intermediate | *ndufs5* | 0.292 | 5.01E-08 |
| 3 | intermediate | *cox4i1* | 0.266 | 7.82E-08 |
| 3 | intermediate | *uqcrc2a* | 0.289 | 2.32E-07 |
| 3 | intermediate | *tpma1* | 0.263 | 3.63E-07 |
| 3 | intermediate | *zgc:101853* | 0.264 | 3.85E-07 |
| 3 | intermediate | *minos1* | 0.258 | 7.14E-07 |
| 3 | intermediate | *si:ch1073-325m22.2* | 0.256 | 8.38E-07 |
| 3 | intermediate | *txlnbb1* | 0.25 | 9.52E-07 |
| 3 | intermediate | *coa6* | 0.285 | 3.78E-06 |
| 3 | intermediate | *cox7a2l* | 0.258 | 5.99E-06 |
| 3 | intermediate | *mdh1ab* | 0.27 | 6.34E-06 |
| 3 | intermediate | *blcap* | 0.266 | 5.40E-05 |
| 4 | fast | *myhc4* | 0.64 | 6.91E-39 |
| 4 | fast | *tnnc21* | 0.41 | 8.01E-29 |
| 4 | fast | *tnni2a.31* | 0.359 | 1.31E-24 |
| 4 | fast | *wu:fj49a02* | 0.69 | 2.62E-24 |
| 4 | fast | *actn3a1* | 0.514 | 8.93E-23 |
| 4 | fast | *XLOC-041970* | 0.765 | 1.02E-22 |
| 4 | fast | *ttn.1* | 0.415 | 2.46E-21 |
| 4 | fast | *si:ch73-367p23.2* | 0.442 | 3.56E-21 |
| 4 | fast | *slc25a41* | 0.374 | 7.36E-21 |
| 4 | fast | *eef2l2* | 0.419 | 1.69E-17 |
| 4 | fast | *atp2a1l1* | 0.326 | 2.46E-16 |
| 4 | fast | *eef1da* | 0.321 | 5.45E-16 |
| 4 | fast | *col1a1b1* | 0.693 | 1.76E-15 |
| 4 | fast | *ryr31* | 0.506 | 2.05E-15 |
| 4 | fast | *tpma2* | 0.346 | 2.12E-15 |
| 4 | fast | *neb* | 0.379 | 3.50E-14 |
| 4 | fast | *ldb3a* | 0.412 | 6.35E-14 |
| 4 | fast | *col11a1b* | 0.451 | 2.39E-13 |
| 4 | fast | *pvalb11* | 0.422 | 5.98E-13 |
| 4 | fast | *col1a1a1* | 0.665 | 7.59E-13 |
| 4 | fast | *pabpc4* | 0.419 | 1.49E-12 |
| 4 | fast | *myoc* | 0.412 | 3.72E-12 |
| 4 | fast | *ttn.2* | 0.329 | 7.91E-12 |
| 4 | fast | *col1a21* | 0.604 | 1.79E-10 |
| 4 | fast | *actn3b* | 0.325 | 2.21E-10 |
| 4 | fast | *thbs2b* | 0.406 | 3.97E-10 |
| 4 | fast | *col12a1a1* | 0.515 | 1.11E-09 |
| 4 | fast | *slc38a3b* | 0.399 | 2.05E-09 |
| 4 | fast | *srl* | 0.353 | 2.28E-09 |
| 4 | fast | *ampd11* | 0.301 | 3.38E-09 |

| cluster | muscle type | marker gene | avg_logFC | p_val_adj (<0.05) |
|---|---|---|---|---|
| 4 | fast | *limch1a* | 0.252 | 9.14E-09 |
| 4 | fast | *smyd1a1* | 0.28 | 9.37E-09 |
| 4 | fast | *adssl1* | 0.327 | 1.69E-08 |
| 4 | fast | *CR354556.3* | 0.39 | 3.60E-08 |
| 4 | fast | *hapln1a* | 0.516 | 4.49E-08 |
| 4 | fast | *myom1a* | 0.391 | 4.55E-08 |
| 4 | fast | *pvalb31* | 0.349 | 9.40E-08 |
| 4 | fast | *hhatla* | 0.312 | 1.73E-07 |
| 4 | fast | *enah* | 0.28 | 4.08E-07 |
| 4 | fast | *mllt11* | 0.386 | 6.16E-07 |
| 4 | fast | *hsc70* | 0.295 | 8.85E-07 |
| 4 | fast | *xirp2a* | 0.378 | 2.20E-06 |
| 4 | fast | *casq1a1* | 0.417 | 2.30E-06 |
| 4 | fast | *rtn4b1* | 0.291 | 3.46E-06 |
| 4 | fast | *bhmt* | 0.295 | 5.98E-06 |
| 4 | fast | *myom2a* | 0.286 | 6.81E-06 |
| 4 | fast | *CABZ01044731.1* | 0.331 | 8.37E-06 |
| 4 | fast | *ryr1b* | 0.331 | 9.68E-06 |
| 4 | fast | *klhl31* | 0.311 | 1.47E-05 |
| 4 | fast | *cacna1sb* | 0.265 | 1.74E-05 |
| 4 | fast | *dcn1* | 0.306 | 1.82E-05 |
| 4 | fast | *abi3bpb* | 0.357 | 3.72E-05 |
| 4 | fast | *desma* | 0.35 | 4.25E-05 |
| 4 | fast | *actc1b1* | 0.258 | 4.41E-05 |
| 4 | fast | *pygma* | 0.281 | 6.21E-05 |
| 4 | fast | *mybpc2b1* | 0.379 | 6.60E-05 |
| 4 | fast | *sparc* | 0.358 | 8.19E-05 |
| 4 | fast | *myog* | 0.289 | 8.98E-05 |
| 4 | fast | *nr4a1* | 0.279 | 0.000125 |
| 4 | fast | *klhl43* | 0.358 | 0.000362 |
| 4 | fast | *synpo2la* | 0.269 | 0.000804 |
| 4 | fast | *unm-hu7910* | 0.259 | 0.000847 |
| 4 | fast | *col12a1b* | 0.303 | 0.000904 |
| 4 | fast | *nrap* | 0.374 | 0.001122 |
| 4 | fast | *abraa* | 0.308 | 0.001207 |
| 4 | fast | *usp13* | 0.278 | 0.001746 |
| 4 | fast | *mef2aa* | 0.526 | 0.001768 |
| 4 | fast | *egfl6* | 0.266 | 0.003709 |
| 4 | fast | *rgcc* | 0.267 | 0.004794 |
| 4 | fast | *tmem182a* | 0.251 | 0.006216 |
| 4 | fast | *si:ch73-43g23.1* | 0.272 | 0.007506 |
| 4 | fast | *CR376766.2* | 0.294 | 0.010854 |

| cluster | muscle type | marker gene | avg_logFC | p_val_adj (<0.05) |
|---|---|---|---|---|
| 4 | fast | *ppp2r3a* | 0.252 | 0.011428 |
| 4 | fast | *serpine1* | 0.262 | 0.021311 |
| 4 | fast | *col5a2a* | 0.252 | 0.028283 |
| 5 | fast | *tppp2* | 0.558 | 9.03E-59 |
| 5 | fast | *stx1b* | 0.604 | 3.01E-55 |
| 5 | fast | *tuba1c* | 1.543 | 9.81E-51 |
| 5 | fast | *crabp1a* | 0.748 | 1.07E-49 |
| 5 | fast | *si:dkey-7j14.5* | 0.576 | 5.93E-48 |
| 5 | fast | *sncga* | 1.554 | 6.67E-48 |
| 5 | fast | *sst6* | 0.61 | 1.15E-47 |
| 5 | fast | *p2rx3a* | 1.364 | 7.70E-46 |
| 5 | fast | *tubb5* | 1.333 | 6.28E-45 |
| 5 | fast | *tuba1a* | 1.721 | 8.35E-45 |
| 5 | fast | *stmn2a* | 1.111 | 1.86E-44 |
| 5 | fast | *ywhag1* | 0.902 | 2.49E-44 |
| 5 | fast | *gnao1a* | 0.606 | 3.66E-44 |
| 5 | fast | *gng3* | 0.6 | 1.41E-43 |
| 5 | fast | *map7d2b* | 0.686 | 7.69E-43 |
| 5 | fast | *elavl3* | 0.688 | 4.42E-42 |
| 5 | fast | *stxbp1a* | 0.734 | 5.02E-42 |
| 5 | fast | *kcnab2a* | 0.829 | 5.89E-41 |
| 5 | fast | *aqp9b* | 1.302 | 7.11E-41 |
| 5 | fast | *si:dkey-46i9.1* | 0.414 | 7.74E-41 |
| 5 | fast | *atp1a3a* | 0.542 | 2.06E-40 |
| 5 | fast | *eno2* | 0.767 | 2.18E-39 |
| 5 | fast | *kif1aa* | 0.509 | 3.66E-39 |
| 5 | fast | *entpd3* | 1.172 | 4.88E-39 |
| 5 | fast | *gnb5a* | 0.489 | 2.95E-38 |
| 5 | fast | *neflb* | 0.563 | 3.16E-38 |
| 5 | fast | *dusp3a* | 0.443 | 3.83E-38 |
| 5 | fast | *vamp1* | 0.728 | 5.44E-38 |
| 5 | fast | *scn8ab* | 0.514 | 6.11E-38 |
| 5 | fast | *prph* | 1.017 | 7.26E-38 |
| 5 | fast | *nmnat2* | 0.416 | 1.13E-37 |
| 5 | fast | *elavl4* | 0.718 | 1.19E-37 |
| 5 | fast | *uchl1* | 1.004 | 1.73E-37 |
| 5 | fast | *zgc:65894* | 0.941 | 1.99E-37 |
| 5 | fast | *map1ab* | 0.608 | 3.03E-37 |
| 5 | fast | *snap25b* | 0.616 | 3.74E-37 |
| 5 | fast | *syt9b* | 0.407 | 4.58E-37 |
| 5 | fast | *map1b* | 0.68 | 9.72E-37 |
| 5 | fast | *calm1b* | 1.136 | 1.17E-36 |

| cluster | muscle type | marker gene | avg_logFC | p_val_adj (<0.05) |
|---|---|---|---|---|
| 5 | fast | *napba* | 0.439 | 1.31E-36 |
| 5 | fast | *cplx2* | 0.842 | 2.05E-36 |
| 5 | fast | *gfap* | 0.928 | 2.82E-36 |
| 5 | fast | *phf24* | 0.471 | 7.73E-36 |
| 5 | fast | *tmsb2* | 0.572 | 1.52E-35 |
| 5 | fast | *atp1a3b* | 0.698 | 1.60E-35 |
| 5 | fast | *zgc:153426* | 0.536 | 1.92E-35 |
| 5 | fast | *stmn2b* | 0.773 | 3.39E-35 |
| 5 | fast | *clip3* | 0.524 | 9.11E-35 |
| 5 | fast | *snap25a* | 1.345 | 9.44E-35 |
| 5 | fast | *spock3* | 0.544 | 1.30E-34 |
| 5 | fast | *pvalb6* | 0.388 | 1.42E-34 |
| 5 | fast | *flj13639* | 0.667 | 1.55E-34 |
| 5 | fast | *ank2a* | 0.942 | 8.11E-34 |
| 5 | fast | *ckbb* | 1.136 | 1.40E-33 |
| 5 | fast | *prnpb* | 0.633 | 2.23E-33 |
| 5 | fast | *sncgb* | 0.516 | 2.23E-33 |
| 5 | fast | *adam22* | 0.331 | 3.27E-33 |
| 5 | fast | *hpca* | 1.398 | 4.90E-33 |
| 5 | fast | *si:dkey-33c12.3* | 0.681 | 5.43E-33 |
| 5 | fast | *kcnt2* | 0.789 | 1.54E-32 |
| 5 | fast | *CABZ01072077.1* | 0.673 | 1.58E-32 |
| 5 | fast | *syngr3b* | 0.553 | 2.38E-32 |
| 5 | fast | *ndrg3a* | 0.832 | 4.05E-32 |
| 5 | fast | *stmn1b* | 1.189 | 8.93E-32 |
| 5 | fast | *p2rx3b* | 0.44 | 1.17E-31 |
| 5 | fast | *cplx2l* | 0.801 | 1.35E-31 |
| 5 | fast | *scn3b* | 0.727 | 2.19E-31 |
| 5 | fast | *sult4a1* | 0.326 | 2.69E-31 |
| 5 | fast | *ndrg4* | 0.567 | 3.81E-31 |
| 5 | fast | *maptb* | 0.516 | 4.95E-31 |
| 5 | fast | *itm2ca* | 0.384 | 8.34E-31 |
| 5 | fast | *ppp3ca* | 0.491 | 1.26E-30 |
| 5 | fast | *inafm2* | 0.684 | 2.86E-30 |
| 5 | fast | *vamp2* | 0.55 | 4.27E-30 |
| 5 | fast | *map4l* | 0.503 | 4.56E-30 |
| 5 | fast | *mbpa* | 1.726 | 9.13E-30 |
| 5 | fast | *clstn2* | 0.384 | 1.09E-29 |
| 5 | fast | *si:ch73-119p20.1* | 0.445 | 4.49E-29 |
| 5 | fast | *sypa* | 0.42 | 5.38E-29 |
| 5 | fast | *rtn1b* | 0.413 | 5.86E-29 |
| 5 | fast | *syt11a* | 0.624 | 8.08E-29 |

| cluster | muscle type | marker gene | avg_logFC | p_val_adj (<0.05) |
|---|---|---|---|---|
| 5 | fast | *trpc1* | 0.384 | 1.96E-28 |
| 5 | fast | *alcamb* | 0.525 | 2.55E-28 |
| 5 | fast | *rufy3* | 0.426 | 2.58E-28 |
| 5 | fast | *XLOC-036173* | 0.369 | 2.98E-28 |
| 5 | fast | *l1camb* | 0.613 | 5.18E-28 |
| 5 | fast | *eml2* | 0.903 | 5.42E-28 |
| 5 | fast | *rusc1* | 0.366 | 7.57E-28 |
| 5 | fast | *cadm3* | 0.385 | 2.10E-27 |
| 5 | fast | *adam23a* | 0.617 | 4.16E-27 |
| 5 | fast | *kif1ab* | 0.515 | 4.54E-27 |
| 5 | fast | *plp1b* | 1.277 | 1.49E-26 |
| 5 | fast | *klc4* | 0.549 | 1.58E-26 |
| 5 | fast | *PCP4* | 0.552 | 6.17E-26 |
| 5 | fast | *flot1a* | 0.508 | 9.35E-26 |
| 5 | fast | *pcp4a* | 0.568 | 9.98E-26 |
| 5 | fast | *adcyap1b* | 0.698 | 1.51E-25 |
| 5 | fast | *pcsk1nl* | 0.596 | 4.26E-25 |
| 5 | fast | *trpc4a* | 0.773 | 7.39E-25 |
| 5 | fast | *stmn4* | 0.429 | 2.87E-24 |
| 5 | fast | *scg3* | 0.51 | 4.38E-24 |
| 5 | fast | *flot2b* | 0.352 | 5.17E-24 |
| 5 | fast | *rab6bb* | 0.699 | 5.36E-24 |
| 5 | fast | *eef1a2* | 0.487 | 6.71E-24 |
| 5 | fast | *nptna* | 0.423 | 7.96E-24 |
| 5 | fast | *scg2a* | 0.441 | 9.54E-24 |
| 5 | fast | *tubb2b* | 0.798 | 3.82E-23 |
| 5 | fast | *cckb* | 1.063 | 5.20E-23 |
| 5 | fast | *si:ch211-113g11.6* | 0.373 | 6.30E-23 |
| 5 | fast | *st8sia5* | 0.334 | 6.53E-23 |
| 5 | fast | *ywhag2* | 0.768 | 5.61E-22 |
| 5 | fast | *calm3a* | 0.717 | 7.08E-22 |
| 5 | fast | *dctn1b* | 0.314 | 2.69E-21 |
| 5 | fast | *anxa13l* | 0.839 | 1.08E-20 |
| 5 | fast | *aldocb* | 0.894 | 3.52E-20 |
| 5 | fast | *fgf13a* | 0.41 | 3.96E-20 |
| 5 | fast | *atp1b2a* | 1.046 | 4.86E-20 |
| 5 | fast | *rab6ba* | 0.422 | 7.39E-20 |
| 5 | fast | *tuba2* | 0.524 | 2.56E-19 |
| 5 | fast | *prkcaa* | 0.416 | 5.70E-18 |
| 5 | fast | *gdi1* | 0.518 | 1.33E-17 |
| 5 | fast | *cd59* | 1.418 | 3.69E-17 |
| 5 | fast | *napbb* | 0.378 | 1.65E-16 |

| cluster | muscle type | marker gene | avg_logFC | p_val_adj (<0.05) |
|---|---|---|---|---|
| 5 | fast | *ngfra* | 0.455 | 1.39E-15 |
| 5 | fast | *cpe* | 0.479 | 1.39E-15 |
| 5 | fast | *fam219ab* | 0.282 | 3.80E-15 |
| 5 | fast | *ribc1* | 0.36 | 4.64E-15 |
| 5 | fast | *nptx1l* | 0.497 | 2.60E-14 |
| 5 | fast | *fkbp1ab* | 0.476 | 4.14E-14 |
| 5 | fast | *pcbp4* | 0.419 | 4.30E-14 |
| 5 | fast | *mllt111* | 0.656 | 1.05E-13 |
| 5 | fast | *hsp90aa1.2* | 0.486 | 1.15E-13 |
| 5 | fast | *dpysl2b* | 0.513 | 1.27E-13 |
| 5 | fast | *aplp1* | 0.405 | 2.39E-13 |
| 5 | fast | *atp6v0cb* | 0.589 | 6.33E-13 |
| 5 | fast | *si:dkey-276j7.1* | 0.47 | 7.89E-13 |
| 5 | fast | *calm1a* | 0.748 | 1.48E-12 |
| 5 | fast | *thy1l* | 0.859 | 2.49E-12 |
| 5 | fast | *kif1b* | 0.512 | 2.56E-12 |
| 5 | fast | *edil3a* | 0.323 | 2.72E-12 |
| 5 | fast | *ank3b* | 0.47 | 3.60E-12 |
| 5 | fast | *XLOC-014441* | 0.315 | 1.10E-11 |
| 5 | fast | *cds2* | 0.271 | 2.62E-11 |
| 5 | fast | *cnp* | 0.361 | 4.84E-11 |
| 5 | fast | *cldn19* | 0.318 | 6.73E-11 |
| 5 | fast | *dpysl3* | 0.449 | 9.09E-11 |
| 5 | fast | *kif21a* | 0.337 | 1.57E-10 |
| 5 | fast | *scdb* | 0.357 | 3.41E-10 |
| 5 | fast | *sptan1* | 0.643 | 4.18E-10 |
| 5 | fast | *calm2b1* | 0.767 | 6.95E-10 |
| 5 | fast | *myoz1b1* | 0.306 | 8.99E-10 |
| 5 | fast | *atp6v1e1b* | 0.427 | 9.22E-10 |
| 5 | fast | *tuba8l3* | 0.391 | 1.01E-09 |
| 5 | fast | *ywhaqa* | 0.395 | 1.65E-09 |
| 5 | fast | *ywhah* | 0.381 | 3.53E-09 |
| 5 | fast | *plcb3* | 0.67 | 3.54E-09 |
| 5 | fast | *ugcg* | 0.279 | 9.32E-09 |
| 5 | fast | *ppp3r1a* | 0.289 | 2.81E-08 |
| 5 | fast | *si:dkeyp-75h12.5* | 0.518 | 3.21E-08 |
| 5 | fast | *gapdhs* | 0.749 | 3.43E-08 |
| 5 | fast | *ywhaba* | 0.523 | 5.06E-08 |
| 5 | fast | *mpz* | 0.874 | 6.71E-08 |
| 5 | fast | *clstn1* | 0.432 | 1.03E-07 |
| 5 | fast | *cfl1* | 0.518 | 1.04E-07 |
| 5 | fast | *gpia* | 0.316 | 1.29E-07 |

| cluster | muscle type | marker gene | avg_logFC | p_val_adj (<0.05) |
|---|---|---|---|---|
| 5 | fast | *hmgb3a* | 0.349 | 2.87E-07 |
| 5 | fast | *palm1b* | 0.328 | 3.13E-07 |
| 5 | fast | *s100b* | 0.633 | 4.80E-07 |
| 5 | fast | *clta* | 0.268 | 1.50E-06 |
| 5 | fast | *clasp2* | 0.317 | 3.43E-06 |
| 5 | fast | *etv5a* | 0.373 | 4.74E-06 |
| 5 | fast | *XLOC-006959* | 0.393 | 6.07E-06 |
| 5 | fast | *actr1* | 0.251 | 6.23E-06 |
| 5 | fast | *serinc1* | 0.473 | 6.24E-06 |
| 5 | fast | *gpm6ab* | 0.333 | 9.47E-06 |
| 5 | fast | *rtn1a* | 0.392 | 1.15E-05 |
| 5 | fast | *cltcb* | 0.345 | 1.70E-05 |
| 5 | fast | *pafah1b1b* | 0.395 | 1.85E-05 |
| 5 | fast | *ccl19a.1* | 0.307 | 1.86E-05 |
| 5 | fast | *rbfox1* | 0.278 | 2.00E-05 |
| 5 | fast | *amd1* | 0.297 | 2.12E-05 |
| 5 | fast | *glrx1* | 0.461 | 2.95E-05 |
| 5 | fast | *fkbp1aa* | 0.365 | 3.31E-05 |
| 5 | fast | *calm3b* | 0.404 | 3.41E-05 |
| 5 | fast | *tspan7b* | 0.285 | 3.68E-05 |
| 5 | fast | *stom* | 0.326 | 5.47E-05 |
| 5 | fast | *mapk3* | 0.382 | 5.50E-05 |
| 5 | fast | *cox6a1l* | 0.437 | 0.000139 |
| 5 | fast | *calm2a* | 0.358 | 0.000223 |
| 5 | fast | *si:ch211-213a13.1* | 0.327 | 0.00026 |
| 5 | fast | *gnb1b* | 0.329 | 0.000339 |
| 5 | fast | *dynlrb1* | 0.261 | 0.000571 |
| 5 | fast | *atp2b2* | 0.318 | 0.000597 |
| 5 | fast | *cd99l2* | 0.346 | 0.000606 |
| 5 | fast | *ywhaqb1* | 0.477 | 0.000654 |
| 5 | fast | *atp6v1aa* | 0.315 | 0.000674 |
| 5 | fast | *ppp2r1ba* | 0.332 | 0.00079 |
| 5 | fast | *tmsb4x1* | 0.489 | 0.000912 |
| 5 | fast | *jpt1b* | 0.314 | 0.000972 |
| 5 | fast | *pfn2l* | 0.33 | 0.001144 |
| 5 | fast | *dnaja2b* | 0.272 | 0.001164 |
| 5 | fast | *cdc42* | 0.336 | 0.001172 |
| 5 | fast | *camk2g1* | 0.431 | 0.002024 |
| 5 | fast | *nudt9* | 0.344 | 0.00222 |
| 5 | fast | *sbf1* | 0.259 | 0.002595 |
| 5 | fast | *CABZ01073795.11* | 0.28 | 0.002837 |
| 5 | fast | *zgc:56493* | 0.342 | 0.003248 |

| cluster | muscle type | marker gene | avg_logFC | p_val_adj (<0.05) |
|---|---|---|---|---|
| 5 | fast | *ly75* | 0.295 | 0.003867 |
| 5 | fast | *vat1* | 0.344 | 0.004318 |
| 5 | fast | *proza* | 0.403 | 0.005247 |
| 5 | fast | *zgc:1533171* | 0.549 | 0.005607 |
| 5 | fast | *skp1* | 0.279 | 0.005901 |
| 5 | fast | *vim* | 0.323 | 0.007464 |
| 5 | fast | *ap2m1a* | 0.285 | 0.008263 |
| 5 | fast | *prnprs3* | 0.311 | 0.012187 |
| 5 | fast | *lamp1b* | 0.265 | 0.012781 |
| 5 | fast | *dynll1* | 0.281 | 0.014237 |
| 5 | fast | *akap12b* | 0.304 | 0.016534 |
| 5 | fast | *scn1ba* | 0.251 | 0.020431 |
| 5 | fast | *pygmb* | 0.253 | 0.02974 |
| 5 | fast | *il6st* | 0.29 | 0.031157 |
| 6 | fast & slow | *mybpc2a* | 1.902 | 8.96E-75 |
| 6 | fast & slow | *tpm1* | 2.838 | 6.75E-63 |
| 6 | fast & slow | *tnni2a.1* | 0.369 | 1.34E-47 |
| 6 | fast & slow | *FO704758.1* | 2.559 | 2.31E-46 |
| 6 | fast & slow | *tbx15* | 0.653 | 2.19E-39 |
| 6 | fast & slow | *tnni2a.2* | 0.723 | 2.09E-37 |
| 6 | fast & slow | *tnni2b.21* | 1.295 | 2.52E-28 |
| 6 | fast & slow | *tcap* | 0.818 | 1.63E-27 |
| 6 | fast & slow | *myoc1* | 0.715 | 7.78E-20 |
| 6 | fast & slow | *atp2a1* | 1.589 | 2.31E-19 |
| 6 | fast & slow | *pvalb7* | 1.342 | 3.25E-19 |
| 6 | fast & slow | *pdlim5a* | 0.649 | 1.76E-18 |
| 6 | fast & slow | *mybpha* | 0.568 | 2.03E-18 |
| 6 | fast & slow | *AGBL1* | 0.319 | 1.04E-17 |
| 6 | fast & slow | *mybpc1* | 1.054 | 2.40E-17 |
| 6 | fast & slow | *btg21* | 0.562 | 6.45E-17 |
| 6 | fast & slow | *mylk4a* | 0.554 | 5.07E-16 |
| 6 | fast & slow | *tpm2* | 0.882 | 1.76E-15 |
| 6 | fast & slow | *clec19a* | 0.303 | 6.55E-15 |
| 6 | fast & slow | *fhl1a* | 0.524 | 7.43E-15 |
| 6 | fast & slow | *pmp22b* | 0.683 | 3.06E-14 |
| 6 | fast & slow | *actn3b1* | 0.438 | 9.89E-14 |
| 6 | fast & slow | *myod1* | 0.57 | 2.23E-13 |
| 6 | fast & slow | *jund1* | 0.534 | 6.97E-13 |
| 6 | fast & slow | *rbp41* | 0.476 | 2.68E-12 |
| 6 | fast & slow | *txnipa1* | 0.435 | 6.83E-12 |
| 6 | fast & slow | *hapln1a1* | 0.552 | 4.11E-11 |
| 6 | fast & slow | *il15l* | 0.341 | 4.46E-11 |

| cluster | muscle type | marker gene | avg_logFC | p_val_adj (<0.05) |
|---|---|---|---|---|
| 6 | fast & slow | *prelid3b* | 0.427 | 1.04E-10 |
| 6 | fast & slow | *nmrk21* | 0.406 | 1.40E-10 |
| 6 | fast & slow | *CABZ01102170.1* | 0.266 | 1.54E-10 |
| 6 | fast & slow | *mustn1b* | 0.368 | 8.88E-10 |
| 6 | fast & slow | *myha* | 2.123 | 9.10E-10 |
| 6 | fast & slow | *smpx* | 0.312 | 1.52E-09 |
| 6 | fast & slow | *ttn.11* | 0.296 | 5.45E-09 |
| 6 | fast & slow | *mpz1* | 0.755 | 7.16E-09 |
| 6 | fast & slow | *si:dkey-261h17.11* | 0.53 | 8.66E-09 |
| 6 | fast & slow | *zbtb17* | 0.534 | 9.91E-09 |
| 6 | fast & slow | *eef2l21* | 0.309 | 1.48E-08 |
| 6 | fast & slow | *htra1b* | 0.348 | 1.50E-08 |
| 6 | fast & slow | *cyr611* | 0.597 | 2.61E-08 |
| 6 | fast & slow | *krt941* | 0.467 | 2.81E-08 |
| 6 | fast & slow | *igfbp1a1* | 0.498 | 4.46E-08 |
| 6 | fast & slow | *col1a1a2* | 0.53 | 7.04E-08 |
| 6 | fast & slow | *tuft1a2* | 0.343 | 1.26E-07 |
| 6 | fast & slow | *col1a1b2* | 0.44 | 1.82E-07 |
| 6 | fast & slow | *flnca* | 0.383 | 2.37E-07 |
| 6 | fast & slow | *synpo2lb* | 0.262 | 3.12E-07 |
| 6 | fast & slow | *hspb1* | 0.486 | 4.15E-07 |
| 6 | fast & slow | *ankrd9* | 0.399 | 4.75E-07 |
| 6 | fast & slow | *ttn.21* | 0.303 | 5.10E-07 |
| 6 | fast & slow | *tnni1d* | 0.781 | 1.05E-06 |
| 6 | fast & slow | *mbpb1* | 0.56 | 1.12E-06 |
| 6 | fast & slow | *marcksl1a1* | 0.326 | 2.49E-06 |
| 6 | fast & slow | *slc20a2* | 0.359 | 2.63E-06 |
| 6 | fast & slow | *nfil3-51* | 0.356 | 4.36E-06 |
| 6 | fast & slow | *nr4a11* | 0.317 | 4.60E-06 |
| 6 | fast & slow | *ldha* | 0.38 | 8.77E-06 |
| 6 | fast & slow | *ryr1a* | 0.318 | 1.09E-05 |
| 6 | fast & slow | *tecra* | 0.259 | 2.15E-05 |
| 6 | fast & slow | *zgc:153981* | 0.303 | 2.15E-05 |
| 6 | fast & slow | *cd81a2* | 0.356 | 2.85E-05 |
| 6 | fast & slow | *myoz2b* | 0.255 | 4.06E-05 |
| 6 | fast & slow | *pdk2b1* | 0.292 | 5.91E-05 |
| 6 | fast & slow | *hdlbpa* | 0.327 | 6.04E-05 |
| 6 | fast & slow | *col1a22* | 0.469 | 8.29E-05 |
| 6 | fast & slow | *igfbp5b1* | 0.426 | 8.77E-05 |
| 6 | fast & slow | *mfap51* | 0.27 | 0.000248 |
| 6 | fast & slow | *srl1* | 0.349 | 0.000364 |
| 6 | fast & slow | *tpm3* | 0.43 | 0.000498 |

| cluster | muscle type | marker gene | avg_logFC | p_val_adj (<0.05) |
|---|---|---|---|---|
| 6 | fast & slow | *dusp11* | 0.315 | 0.00052 |
| 6 | fast & slow | *txnipb* | 0.286 | 0.000551 |
| 6 | fast & slow | *ppdpfa* | 0.29 | 0.000581 |
| 6 | fast & slow | *dnajb6b* | 0.265 | 0.000847 |
| 6 | fast & slow | *col12a1a2* | 0.372 | 0.000953 |
| 6 | fast & slow | *XLOC-027924* | 0.303 | 0.001531 |
| 6 | fast & slow | *thbs3a* | 0.262 | 0.001751 |
| 6 | fast & slow | *dcn2* | 0.376 | 0.002547 |
| 6 | fast & slow | *clu2* | 0.293 | 0.003682 |
| 6 | fast & slow | *gyg1a* | 0.314 | 0.004157 |
| 6 | fast & slow | *rtn3* | 0.3 | 0.004417 |
| 6 | fast & slow | *si:ch211-5k11.8* | 0.369 | 0.005415 |
| 6 | fast & slow | *clic4* | 0.327 | 0.00559 |
| 6 | fast & slow | *mibp* | 0.273 | 0.005652 |
| 6 | fast & slow | *gapdh1* | 0.32 | 0.006065 |
| 6 | fast & slow | *CABZ01072309.1* | 0.324 | 0.006678 |
| 6 | fast & slow | *tmem176l.1* | 0.288 | 0.006873 |
| 6 | fast & slow | *creg1* | 0.262 | 0.006875 |
| 6 | fast & slow | *g0s2* | 0.379 | 0.008573 |
| 6 | fast & slow | *crip12* | 0.282 | 0.009111 |
| 6 | fast & slow | *klhl21* | 0.289 | 0.009829 |
| 6 | fast & slow | *hspg2* | 0.274 | 0.012093 |
| 6 | fast & slow | *thbs1a* | 0.376 | 0.012375 |
| 6 | fast & slow | *zgc:112356* | 0.268 | 0.023542 |
| 6 | fast & slow | *desma1* | 0.39 | 0.026682 |
| 6 | fast & slow | *serbp1b* | 0.304 | 0.029517 |
| 6 | fast & slow | *ehbp1l1a* | 0.254 | 0.029848 |
| 6 | fast & slow | *sgcg* | 0.291 | 0.039341 |
| 6 | fast & slow | *zfand5b* | 0.281 | 0.040642 |
| 7 | slow | *tnni4b.2* | 1.178 | ####### |
| 7 | slow | *tnni4b.1* | 1.675 | 2.37E-93 |
| 7 | slow | *mybpc3* | 1.343 | 6.63E-84 |
| 7 | slow | *smyhc2* | 2.109 | 4.67E-78 |
| 7 | slow | *cox4i1l* | 1.568 | 1.09E-72 |
| 7 | slow | *myoz2b1* | 0.692 | 3.72E-72 |
| 7 | slow | *CU633479.5* | 0.971 | 2.33E-71 |
| 7 | slow | *tnnc1b* | 2.149 | 3.44E-71 |
| 7 | slow | *LOC110440135* | 0.683 | 1.85E-64 |
| 7 | slow | *actc1c* | 2.727 | 1.17E-62 |
| 7 | slow | *tnnt2e* | 2.062 | 4.43E-62 |
| 7 | slow | *tnni1c* | 2.245 | 6.40E-62 |
| 7 | slow | *myl10* | 2.837 | 1.78E-61 |

| cluster | muscle type | marker gene | avg_logFC | p_val_adj (<0.05) |
|---|---|---|---|---|
| 7 | slow | *tnnt2d* | 0.727 | 5.93E-60 |
| 7 | slow | *myl13* | 2.624 | 1.24E-58 |
| 7 | slow | *slc25a51* | 1.59 | 3.18E-56 |
| 7 | slow | *COX31* | 1.251 | 1.35E-55 |
| 7 | slow | *mt-cyb1* | 1.508 | 4.34E-55 |
| 7 | slow | *mt-co21* | 1.201 | 8.93E-55 |
| 7 | slow | *tpm21* | 1.947 | 2.15E-54 |
| 7 | slow | *mustn1b1* | 0.744 | 3.22E-53 |
| 7 | slow | *NC-002333.171* | 1.293 | 7.65E-53 |
| 7 | slow | *casq2* | 0.533 | 1.42E-52 |
| 7 | slow | *ND41* | 1.316 | 6.06E-52 |
| 7 | slow | *atp2a2a* | 2.58 | 1.76E-51 |
| 7 | slow | *ATP61* | 1.458 | 1.14E-50 |
| 7 | slow | *mt-co11* | 1.192 | 4.76E-50 |
| 7 | slow | *mt-nd11* | 1.276 | 1.37E-49 |
| 7 | slow | *ND31* | 1.322 | 2.83E-49 |
| 7 | slow | *atp5f1b* | 0.981 | 3.22E-49 |
| 7 | slow | *pvalb71* | 2.002 | 4.01E-49 |
| 7 | slow | *trnS21* | 0.798 | 8.01E-49 |
| 7 | slow | *atp5mc3b1* | 0.767 | 1.60E-47 |
| 7 | slow | *atp5fa1* | 0.982 | 1.03E-46 |
| 7 | slow | *tnni1d1* | 0.928 | 1.66E-46 |
| 7 | slow | *ND21* | 1.192 | 1.09E-45 |
| 7 | slow | *hspb6* | 1.033 | 1.33E-44 |
| 7 | slow | *hhatlb* | 0.475 | 1.71E-44 |
| 7 | slow | *cox6a21* | 1.114 | 2.98E-44 |
| 7 | slow | *idh21* | 0.713 | 9.68E-44 |
| 7 | slow | *si:ch73-288o11.5* | 0.858 | 1.29E-43 |
| 7 | slow | *atp5mc11* | 0.921 | 2.10E-43 |
| 7 | slow | *ryr1a1* | 0.585 | 6.16E-43 |
| 7 | slow | *coq8a* | 0.797 | 8.22E-43 |
| 7 | slow | *atp2a11* | 1.325 | 3.27E-42 |
| 7 | slow | *mdh1aa* | 1.141 | 3.81E-41 |
| 7 | slow | *mb* | 1.382 | 4.16E-41 |
| 7 | slow | *LOC100535869* | 0.328 | 5.96E-41 |
| 7 | slow | *fabp3* | 1.139 | 1.63E-40 |
| 7 | slow | *tnni2a.41* | 1.779 | 1.08E-37 |
| 7 | slow | *BX248497.1* | 0.276 | 2.47E-36 |
| 7 | slow | *NC-002333.42* | 0.634 | 3.95E-36 |
| 7 | slow | *cox8b* | 1.088 | 6.01E-36 |
| 7 | slow | *si:ch1073-325m22.21* | 0.881 | 7.51E-36 |
| 7 | slow | *CABZ01076275.1* | 0.97 | 1.85E-35 |

| cluster | muscle type | marker gene | avg_logFC | p_val_adj (<0.05) |
|---|---|---|---|---|
| 7 | slow | *LO018250.1* | 0.354 | 1.91E-35 |
| 7 | slow | *mdh21* | 0.648 | 1.37E-34 |
| 7 | slow | *got2a* | 0.874 | 3.12E-34 |
| 7 | slow | *rnf207b* | 0.375 | 1.05E-33 |
| 7 | slow | *tpm31* | 1.67 | 3.96E-33 |
| 7 | slow | *atp5f1c1* | 0.724 | 1.96E-32 |
| 7 | slow | *tnni4a* | 0.545 | 1.68E-31 |
| 7 | slow | *lpl* | 0.765 | 5.72E-31 |
| 7 | slow | *smpx1* | 0.477 | 1.56E-30 |
| 7 | slow | *ndufa4l1* | 0.764 | 2.43E-30 |
| 7 | slow | *slc25a3b* | 0.58 | 3.93E-30 |
| 7 | slow | *sdha* | 0.805 | 4.89E-30 |
| 7 | slow | *smyhc3* | 0.601 | 9.16E-30 |
| 7 | slow | *NC-002333.141* | 0.746 | 1.05E-29 |
| 7 | slow | *cox6c1* | 0.736 | 1.36E-29 |
| 7 | slow | *uqcrfs1* | 0.78 | 3.74E-29 |
| 7 | slow | *cox5b21* | 0.86 | 2.99E-28 |
| 7 | slow | *atp5pf1* | 0.705 | 3.99E-28 |
| 7 | slow | *hadhab* | 0.551 | 6.90E-28 |
| 7 | slow | *uqcrc2b* | 0.696 | 1.41E-27 |
| 7 | slow | *cox5ab* | 0.807 | 1.43E-27 |
| 7 | slow | *uqcrc1* | 0.752 | 1.59E-27 |
| 7 | slow | *cox7b* | 0.825 | 1.59E-27 |
| 7 | slow | *bckdha* | 0.416 | 1.64E-27 |
| 7 | slow | *CU633479.2* | 0.581 | 5.04E-27 |
| 7 | slow | *uqcrh* | 0.759 | 8.74E-27 |
| 7 | slow | *tcap1* | 0.61 | 2.52E-26 |
| 7 | slow | *psd3l* | 0.418 | 1.07E-25 |
| 7 | slow | *acadvl* | 0.776 | 1.18E-25 |
| 7 | slow | *cycsb* | 0.723 | 1.47E-25 |
| 7 | slow | *ndufb71* | 0.68 | 2.70E-25 |
| 7 | slow | *cpt1b* | 0.352 | 4.55E-25 |
| 7 | slow | *prodha* | 0.561 | 1.75E-24 |
| 7 | slow | *atp5mf* | 0.673 | 1.84E-24 |
| 7 | slow | *atp5meb1* | 0.546 | 3.23E-24 |
| 7 | slow | *atp5pd1* | 0.653 | 5.70E-24 |
| 7 | slow | *atp5pb1* | 0.644 | 6.07E-24 |
| 7 | slow | *hbba1* | 0.887 | 7.84E-24 |
| 7 | slow | *zgc:1935411* | 0.57 | 7.85E-24 |
| 7 | slow | *atp5po1* | 0.606 | 8.89E-24 |
| 7 | slow | *ndufs7* | 0.784 | 1.16E-23 |
| 7 | slow | *acat1* | 0.611 | 1.37E-23 |

| cluster | muscle type | marker gene | avg_logFC | p_val_adj (<0.05) |
|---|---|---|---|---|
| 7 | slow | *si:ch211-5k11.81* | 0.907 | 1.55E-23 |
| 7 | slow | *cox7c1* | 0.705 | 1.93E-23 |
| 7 | slow | *aco2* | 0.786 | 2.05E-23 |
| 7 | slow | *oxct1a* | 0.583 | 2.82E-23 |
| 7 | slow | *atp5if1b* | 0.78 | 4.10E-23 |
| 7 | slow | *cox4i1l* | 0.708 | 5.09E-23 |
| 7 | slow | *ldhba* | 0.962 | 9.01E-23 |
| 7 | slow | *etfa* | 0.692 | 1.11E-22 |
| 7 | slow | *got1* | 0.623 | 1.53E-22 |
| 7 | slow | *esrra* | 0.457 | 1.64E-22 |
| 7 | slow | *si:ch73-390p7.2* | 0.312 | 1.91E-22 |
| 7 | slow | *pdha1a* | 0.673 | 2.26E-22 |
| 7 | slow | *ckmt2b1* | 0.697 | 3.42E-22 |
| 7 | slow | *pfkmb* | 0.627 | 2.29E-21 |
| 7 | slow | *acadm* | 0.724 | 2.95E-21 |
| 7 | slow | *ndufab1b* | 0.652 | 3.16E-21 |
| 7 | slow | *cyc1* | 0.716 | 3.55E-21 |
| 7 | slow | *atp5md1* | 0.675 | 3.90E-21 |
| 7 | slow | *coq10b1* | 0.613 | 5.62E-21 |
| 7 | slow | *cs* | 0.633 | 6.25E-21 |
| 7 | slow | *mdh1ab1* | 0.595 | 1.44E-20 |
| 7 | slow | *atp5f1d1* | 0.591 | 2.99E-20 |
| 7 | slow | *cox7a1l* | 0.667 | 4.17E-20 |
| 7 | slow | *ppdpfa1* | 0.608 | 4.99E-20 |
| 7 | slow | *mpc1l* | 0.644 | 9.71E-20 |
| 7 | slow | *ndufb8* | 0.604 | 1.78E-19 |
| 7 | slow | *zgc:85777* | 0.612 | 1.80E-19 |
| 7 | slow | *cox6b1* | 0.482 | 5.72E-19 |
| 7 | slow | *gpd1a* | 0.306 | 1.04E-18 |
| 7 | slow | *XLOC-032994* | 0.325 | 2.78E-18 |
| 7 | slow | *cox6b21* | 0.557 | 3.11E-18 |
| 7 | slow | *ndufs3* | 0.649 | 6.43E-18 |
| 7 | slow | *hbaa1l* | 0.657 | 1.28E-17 |
| 7 | slow | *hsd17b10* | 0.301 | 2.31E-17 |
| 7 | slow | *ndufb2* | 0.659 | 3.80E-17 |
| 7 | slow | *chchd6b* | 0.535 | 3.87E-17 |
| 7 | slow | *lyrm7* | 0.345 | 6.81E-17 |
| 7 | slow | *LOC100331743* | 0.534 | 1.02E-16 |
| 7 | slow | *flnca1* | 0.531 | 1.16E-16 |
| 7 | slow | *cdh5* | 0.352 | 1.30E-16 |
| 7 | slow | *dlat* | 0.573 | 1.84E-16 |
| 7 | slow | *tnnt1* | 0.499 | 2.20E-16 |

| cluster | muscle type | marker gene | avg_logFC | p_val_adj (<0.05) |
|---|---|---|---|---|
| 7 | slow | *ndufs8a* | 0.49 | 6.06E-16 |
| 7 | slow | *chchd10* | 0.623 | 8.47E-16 |
| 7 | slow | *zbtb171* | 0.394 | 8.95E-16 |
| 7 | slow | *hsdl2* | 0.374 | 1.13E-15 |
| 7 | slow | *eci1* | 0.527 | 1.99E-15 |
| 7 | slow | *atp5l1* | 0.466 | 2.04E-15 |
| 7 | slow | *atp5mc3a* | 0.654 | 2.29E-15 |
| 7 | slow | *ndufa10* | 0.537 | 3.34E-15 |
| 7 | slow | *atp1b1a2* | 0.48 | 3.40E-15 |
| 7 | slow | *pptc7b* | 0.422 | 3.65E-15 |
| 7 | slow | *mybpc11* | 0.709 | 4.72E-15 |
| 7 | slow | *acta1b* | 0.267 | 5.48E-15 |
| 7 | slow | *ndufab1a* | 0.558 | 6.06E-15 |
| 7 | slow | *slc16a1a* | 0.526 | 8.51E-15 |
| 7 | slow | *ogdha* | 0.519 | 1.15E-14 |
| 7 | slow | *mpc2* | 0.479 | 2.23E-14 |
| 7 | slow | *lbh* | 0.343 | 7.39E-14 |
| 7 | slow | *etfdh* | 0.366 | 1.06E-13 |
| 7 | slow | *fabp11a* | 0.601 | 1.44E-13 |
| 7 | slow | *cpt1ab* | 0.326 | 2.06E-13 |
| 7 | slow | *echs1* | 0.348 | 4.03E-13 |
| 7 | slow | *ndufa8* | 0.475 | 4.37E-13 |
| 7 | slow | *acads* | 0.293 | 7.68E-13 |
| 7 | slow | *si:ch211-171h4.3* | 0.26 | 7.89E-13 |
| 7 | slow | *fhl1a1* | 0.368 | 8.20E-13 |
| 7 | slow | *NDUFB11* | 0.522 | 8.64E-13 |
| 7 | slow | *slc16a1b* | 0.409 | 9.07E-13 |
| 7 | slow | *ndufa6* | 0.447 | 1.03E-12 |
| 7 | slow | *mrps36* | 0.434 | 1.06E-12 |
| 7 | slow | *synpo2lb1* | 0.361 | 1.11E-12 |
| 7 | slow | *mt-nd51* | 0.488 | 1.25E-12 |
| 7 | slow | *dldh* | 0.509 | 1.39E-12 |
| 7 | slow | *trnR* | 0.292 | 1.55E-12 |
| 7 | slow | *vegfab* | 0.314 | 1.80E-12 |
| 7 | slow | *vdac3* | 0.478 | 1.97E-12 |
| 7 | slow | *sod2* | 0.46 | 2.23E-12 |
| 7 | slow | *bckdhb* | 0.315 | 2.29E-12 |
| 7 | slow | *glud1b* | 0.514 | 2.68E-12 |
| 7 | slow | *mibp1* | 0.485 | 2.99E-12 |
| 7 | slow | *lyar* | 0.328 | 3.55E-12 |
| 7 | slow | *fstl1a* | 0.275 | 3.80E-12 |
| 7 | slow | *sdhb* | 0.476 | 4.08E-12 |

| cluster | muscle type | marker gene | avg_logFC | p_val_adj (<0.05) |
|---|---|---|---|---|
| 7 | slow | *acsl1b* | 0.384 | 4.12E-12 |
| 7 | slow | *bpgm* | 0.34 | 4.28E-12 |
| 7 | slow | *acadl* | 0.395 | 4.52E-12 |
| 7 | slow | *hadhaa* | 0.346 | 4.58E-12 |
| 7 | slow | *pfkfb2b* | 0.252 | 5.01E-12 |
| 7 | slow | *hadhb* | 0.531 | 5.90E-12 |
| 7 | slow | *ivd* | 0.345 | 7.03E-12 |
| 7 | slow | *nipsnap2* | 0.464 | 7.89E-12 |
| 7 | slow | *CABZ01084963.1* | 0.461 | 8.59E-12 |
| 7 | slow | *mut* | 0.449 | 1.00E-11 |
| 7 | slow | *rgs5a1* | 0.433 | 1.34E-11 |
| 7 | slow | *uqcr101* | 0.487 | 1.58E-11 |
| 7 | slow | *alas11* | 0.422 | 2.93E-11 |
| 7 | slow | *prdx3* | 0.438 | 3.43E-11 |
| 7 | slow | *ndufa5* | 0.525 | 4.66E-11 |
| 7 | slow | *mfsd2ab1* | 0.637 | 4.67E-11 |
| 7 | slow | *tnni2b.22* | 0.649 | 4.96E-11 |
| 7 | slow | *bckdk* | 0.398 | 6.75E-11 |
| 7 | slow | *si:dkey-98j1.51* | 0.466 | 1.26E-10 |
| 7 | slow | *arg21* | 0.471 | 1.50E-10 |
| 7 | slow | *ndufa9a* | 0.505 | 1.58E-10 |
| 7 | slow | *ndufc2* | 0.494 | 1.68E-10 |
| 7 | slow | *pgk1* | 0.439 | 2.70E-10 |
| 7 | slow | *bsg1* | 0.469 | 3.24E-10 |
| 7 | slow | *ndufb10* | 0.429 | 3.31E-10 |
| 7 | slow | *sucla2* | 0.418 | 3.38E-10 |
| 7 | slow | *uqcrq1* | 0.487 | 3.75E-10 |
| 7 | slow | *immt* | 0.385 | 3.90E-10 |
| 7 | slow | *fxyd11* | 0.338 | 4.32E-10 |
| 7 | slow | *sdhc* | 0.512 | 4.80E-10 |
| 7 | slow | *fth1a1* | 0.343 | 5.68E-10 |
| 7 | slow | *CABZ01072309.11* | 0.477 | 5.94E-10 |
| 7 | slow | *myoz3a* | 0.34 | 6.24E-10 |
| 7 | slow | *mrpl47* | 0.304 | 7.59E-10 |
| 7 | slow | *ndufv2* | 0.397 | 1.21E-09 |
| 7 | slow | *etfb* | 0.406 | 1.24E-09 |
| 7 | slow | *ndufs6* | 0.463 | 1.31E-09 |
| 7 | slow | *camk2a* | 0.263 | 1.52E-09 |
| 7 | slow | *slc20a21* | 0.401 | 1.72E-09 |
| 7 | slow | *atp5if1a1* | 0.498 | 1.91E-09 |
| 7 | slow | *dlst* | 0.487 | 2.11E-09 |
| 7 | slow | *hspb12* | 0.517 | 2.46E-09 |

| cluster | muscle type | marker gene | avg_logFC | p_val_adj (<0.05) |
|---|---|---|---|---|
| 7 | slow | *tpi1b1* | 0.384 | 3.38E-09 |
| 7 | slow | *suclg1* | 0.457 | 4.21E-09 |
| 7 | slow | *ndufs51* | 0.414 | 7.17E-09 |
| 7 | slow | *slc16a3* | 0.315 | 9.09E-09 |
| 7 | slow | *hibadhb* | 0.251 | 9.67E-09 |
| 7 | slow | *acad8* | 0.264 | 1.08E-08 |
| 7 | slow | *uqcrc2a1* | 0.436 | 1.36E-08 |
| 7 | slow | *mlip* | 0.391 | 1.43E-08 |
| 7 | slow | *si:ch211-235e9.6* | 0.395 | 1.54E-08 |
| 7 | slow | *ugp2a* | 0.374 | 2.02E-08 |
| 7 | slow | *aifm1* | 0.329 | 2.31E-08 |
| 7 | slow | *aqp8a.1* | 0.444 | 2.58E-08 |
| 7 | slow | *apooa* | 0.316 | 2.88E-08 |
| 7 | slow | *ndufa12* | 0.405 | 2.90E-08 |
| 7 | slow | *mccc1* | 0.256 | 2.99E-08 |
| 7 | slow | *fhl2a* | 0.301 | 3.80E-08 |
| 7 | slow | *XLOC-0196721* | 0.363 | 3.91E-08 |
| 7 | slow | *cox5aa* | 0.389 | 4.47E-08 |
| 7 | slow | *ndufb6* | 0.415 | 4.50E-08 |
| 7 | slow | *hspd1* | 0.27 | 5.27E-08 |
| 7 | slow | *zfand5b1* | 0.365 | 5.86E-08 |
| 7 | slow | *ppp1r3ca* | 0.273 | 6.11E-08 |
| 7 | slow | *mybpha1* | 0.278 | 6.37E-08 |
| 7 | slow | *ndufv3* | 0.416 | 7.68E-08 |
| 7 | slow | *gcat* | 0.288 | 1.20E-07 |
| 7 | slow | *glula* | 0.306 | 1.28E-07 |
| 7 | slow | *cavin1b* | 0.349 | 1.32E-07 |
| 7 | slow | *ndufs41* | 0.384 | 1.33E-07 |
| 7 | slow | *ndufa21* | 0.453 | 1.92E-07 |
| 7 | slow | *rtn1a1* | 0.343 | 2.21E-07 |
| 7 | slow | *coq9* | 0.363 | 2.36E-07 |
| 7 | slow | *nmrk22* | 0.348 | 2.70E-07 |
| 7 | slow | *sdhdb* | 0.353 | 3.84E-07 |
| 7 | slow | *ndufs2* | 0.401 | 4.30E-07 |
| 7 | slow | *si:zfos-1192g2.3* | 0.307 | 5.16E-07 |
| 7 | slow | *aldh6a1* | 0.389 | 6.88E-07 |
| 7 | slow | *slc25a33* | 0.374 | 7.14E-07 |
| 7 | slow | *stau2* | 0.351 | 7.39E-07 |
| 7 | slow | *gstr* | 0.377 | 1.12E-06 |
| 7 | slow | *pdk2b2* | 0.332 | 1.16E-06 |
| 7 | slow | *oat* | 0.419 | 1.22E-06 |
| 7 | slow | *pdhb* | 0.507 | 1.30E-06 |

| cluster | muscle type | marker gene | avg_logFC | p_val_adj (<0.05) |
|---|---|---|---|---|
| 7 | slow | *gatd3a* | 0.275 | 2.27E-06 |
| 7 | slow | *slc6a16a* | 0.329 | 2.36E-06 |
| 7 | slow | *ppifb* | 0.313 | 2.46E-06 |
| 7 | slow | *mfn2* | 0.287 | 2.58E-06 |
| 7 | slow | *gcdha* | 0.394 | 2.94E-06 |
| 7 | slow | *eno32* | 0.335 | 3.87E-06 |
| 7 | slow | *cav1* | 0.303 | 3.87E-06 |
| 7 | slow | *mylk4a1* | 0.353 | 4.77E-06 |
| 7 | slow | *jund2* | 0.306 | 5.64E-06 |
| 7 | slow | *eci2* | 0.252 | 8.56E-06 |
| 7 | slow | *zgc:162509* | 0.267 | 1.11E-05 |
| 7 | slow | *desma2* | 0.458 | 1.21E-05 |
| 7 | slow | *ndufb9* | 0.37 | 1.49E-05 |
| 7 | slow | *cox7a2l1* | 0.314 | 1.66E-05 |
| 7 | slow | *sbk3* | 0.257 | 1.69E-05 |
| 7 | slow | *arl4ab* | 0.291 | 2.97E-05 |
| 7 | slow | *slc25a111* | 0.373 | 3.06E-05 |
| 7 | slow | *si:ch211-250g4.3* | 0.295 | 3.13E-05 |
| 7 | slow | *igfbp7* | 0.26 | 3.47E-05 |
| 7 | slow | *ndufa13* | 0.32 | 4.10E-05 |
| 7 | slow | *mybpc2b2* | 0.456 | 4.26E-05 |
| 7 | slow | *hspb111* | 0.446 | 4.27E-05 |
| 7 | slow | *hspb8* | 0.27 | 6.50E-05 |
| 7 | slow | *slc25a55a2* | 0.258 | 7.18E-05 |
| 7 | slow | *eif4a1b* | 0.28 | 8.61E-05 |
| 7 | slow | *pygmb1* | 0.288 | 0.000127 |
| 7 | slow | *si:ch1073-314i13.4* | 0.346 | 0.000147 |
| 7 | slow | *pdhx* | 0.293 | 0.000167 |
| 7 | slow | *got2b* | 0.325 | 0.000169 |
| 7 | slow | *prelid3b1* | 0.31 | 0.00017 |
| 7 | slow | *me3* | 0.345 | 0.00025 |
| 7 | slow | *atp5f1e2* | 0.272 | 0.00025 |
| 7 | slow | *srl2* | 0.28 | 0.000347 |
| 7 | slow | *hadh* | 0.283 | 0.000384 |
| 7 | slow | *ndufb41* | 0.308 | 0.000415 |
| 7 | slow | *ndufb5* | 0.338 | 0.000455 |
| 7 | slow | *cox7a2a* | 0.256 | 0.000584 |
| 7 | slow | *ndufv1* | 0.344 | 0.000664 |
| 7 | slow | *klhl431* | 0.308 | 0.000732 |
| 7 | slow | *smyd1b* | 0.318 | 0.000756 |
| 7 | slow | *cregl1* | 0.305 | 0.001022 |
| 7 | slow | *slc25a12* | 0.316 | 0.001212 |

| cluster | muscle type | marker gene | avg_logFC | p_val_adj (<0.05) |
|---|---|---|---|---|
| 7 | slow | *atp5mea* | 0.308 | 0.001269 |
| 7 | slow | *slc7a1* | 0.274 | 0.001515 |
| 7 | slow | *fh1* | 0.331 | 0.001666 |
| 7 | slow | *XLOC-001733* | 0.311 | 0.001876 |
| 7 | slow | *vdac21* | 0.283 | 0.00189 |
| 7 | slow | *gys1* | 0.328 | 0.002466 |
| 7 | slow | *crip13* | 0.313 | 0.003729 |
| 7 | slow | *nfil3-62* | 0.267 | 0.003951 |
| 7 | slow | *sorbs11* | 0.254 | 0.004041 |
| 7 | slow | *ndufa11* | 0.331 | 0.004502 |
| 7 | slow | *hspe1* | 0.289 | 0.005409 |
| 7 | slow | *qki2* | 0.256 | 0.006253 |
| 7 | slow | *phb* | 0.26 | 0.007268 |
| 7 | slow | *hspa9* | 0.278 | 0.007872 |
| 7 | slow | *minos11* | 0.272 | 0.015943 |
| 7 | slow | *XLOC-022132* | 0.25 | 0.016685 |
| 7 | slow | *chchd2* | 0.272 | 0.021632 |
| 7 | slow | *ndufa31* | 0.32 | 0.025689 |
| 7 | slow | *ndufa71* | 0.299 | 0.030933 |
| 7 | slow | *ckmt2a1* | 0.281 | 0.039143 |
| 7 | slow | *srgn* | 0.261 | 0.049563 |
| 8 | | *hbaa12* | 1.859 | 2.80E-31 |
| 8 | | *hbba11* | 1.501 | 1.33E-26 |
| 8 | | *si:ch211-5k11.82* | 1.134 | 4.25E-20 |
| 8 | | *si:ch211-250g4.31* | 1.085 | 1.67E-17 |
| 8 | | *mybphb1* | 0.593 | 1.94E-12 |
| 8 | | *si:dkey-151g10.61* | 0.332 | 2.89E-09 |
| 8 | | *tnnt3b1* | 0.299 | 9.45E-08 |
| 8 | | *ckma2* | 0.322 | 1.14E-07 |
| 8 | | *uqcrb2* | 0.412 | 1.14E-07 |
| 8 | | *si:dkey-211f22.51* | 0.423 | 1.93E-07 |
| 8 | | *myl11* | 0.393 | 8.69E-07 |
| 8 | | *si:dkey-51e6.11* | 0.362 | 9.51E-07 |
| 8 | | *pvalb21* | 0.475 | 1.22E-06 |
| 8 | | *aldoab2* | 0.317 | 4.11E-06 |
| 8 | | *aldoaa2* | 0.32 | 5.09E-06 |
| 8 | | *rpl381* | 0.311 | 6.03E-06 |
| 8 | | *mylpfa1* | 0.278 | 1.59E-05 |
| 8 | | *tpma3* | 0.303 | 2.82E-05 |
| 8 | | *ampd12* | 0.337 | 6.07E-05 |
| 8 | | *rpl27a1* | 0.293 | 0.000129 |
| 8 | | *ckmb2* | 0.25 | 0.000148 |

| cluster | muscle type | marker gene | avg_logFC | p_val_adj (<0.05) |
|---|---|---|---|---|
| 8 | | *atp5f1e3* | 0.297 | 0.000149 |
| 8 | | *actn3a2* | 0.343 | 0.000156 |
| 8 | | *CABZ01078594.11* | 0.359 | 0.000214 |
| 8 | | *tmod41* | 0.33 | 0.000224 |
| 8 | | *smyd1a2* | 0.29 | 0.000631 |
| 8 | | *ppp1cbl* | 0.312 | 0.001865 |
| 8 | | *mylz31* | 0.267 | 0.002329 |
| 8 | | *mkrn1* | 0.26 | 0.007155 |
| 8 | | *pkmb1* | 0.251 | 0.013383 |
| 8 | | *pvalb32* | 0.268 | 0.013986 |
| 8 | | *bhmt1* | 0.273 | 0.01825 |
| 8 | | *zgc:1018531* | 0.265 | 0.018903 |
| 8 | | *txlnbb2* | 0.28 | 0.024323 |
| 8 | | *si:ch211-255p10.31* | 0.261 | 0.034332 |
| 8 | | *rpl91* | 0.281 | 0.047635 |
| 9 | | *sf3b6* | 0.362 | 6.61E-05 |
| 9 | | *pvalb22* | 0.75 | 0.000106 |
| 9 | | *kbtbd12* | 0.457 | 0.00011 |
| 9 | | *scn4ab* | 0.453 | 0.00402 |
| 9 | | *pvalb13* | 0.468 | 0.008766 |
| 9 | | *six1a* | 0.291 | 0.016353 |
