## Supplementary material for "Zebrafish *Danio rerio* trunk muscle structure and growth from a spatial transcriptomics perspective": Marker genes of each cluster in larvae zebrafish muscle

supplementary 3 Marker genes of each cluster in larvae zebrafish (p<0.05)

| cluster | marker gene | avg_logFC | p_val_adj (<0.05) |
|---|---|---|---|
| 0 | *krt17* | 1.496774 | 5.51E-12 |
| 0 | *actb2* | 0.916902 | 7.10E-12 |
| 0 | *icn* | 1.215925 | 1.31E-11 |
| 0 | *krt91* | 1.33164 | 2.82E-11 |
| 0 | *krt4* | 1.336794 | 3.71E-11 |
| 0 | *krt5* | 1.263112 | 4.35E-11 |
| 0 | *tmsb4x* | 0.791429 | 1.51E-10 |
| 0 | *si:dkey-7c18.24* | 1.17412 | 2.86E-09 |
| 0 | *mcl1a* | 1.009817 | 3.84E-09 |
| 0 | *zgc:111983* | 1.223962 | 4.74E-09 |
| 0 | *icn2* | 1.130969 | 5.07E-09 |
| 0 | *actb1* | 0.96943 | 8.67E-09 |
| 0 | *tmsb1* | 0.932385 | 1.14E-08 |
| 0 | *si:dkey-248g15.3* | 1.664966 | 1.63E-07 |
| 0 | *zgc:193505* | 1.199538 | 1.74E-07 |
| 0 | *klf2a* | 1.111599 | 1.79E-07 |
| 0 | *cfl1l* | 1.014567 | 1.99E-07 |
| 0 | *s100a10b* | 0.815137 | 2.09E-07 |
| 0 | *pfn1* | 0.890315 | 2.33E-07 |
| 0 | *wu:fb18f06* | 0.996333 | 2.77E-07 |
| 0 | *anxa2a* | 0.802437 | 8.84E-07 |
| 0 | *ppiaa* | 0.539967 | 8.88E-07 |
| 0 | *anxa1a* | 0.975534 | 9.29E-07 |
| 0 | *cldnb* | 1.044908 | 1.55E-06 |
| 0 | *ptgdsb.1* | 1.126965 | 1.59E-06 |
| 0 | *apoeb* | 1.142415 | 2.32E-06 |
| 0 | *si:dkey-193p11.2* | 1.256373 | 3.15E-06 |
| 0 | *ptgdsb.2* | 0.990913 | 7.21E-06 |
| 0 | *cldni* | 0.8949 | 1.32E-05 |
| 0 | *mhc1zba* | 0.929971 | 1.71E-05 |
| 0 | *epcam* | 0.939377 | 1.95E-05 |
| 0 | *jupa* | 0.819298 | 2.95E-05 |
| 0 | *dsg2.1* | 0.880573 | 3.69E-05 |
| 0 | *cfd* | 0.862903 | 3.94E-05 |
| 0 | *anxa1c* | 0.745864 | 4.26E-05 |
| 0 | *bzw1b* | 0.923753 | 4.75E-05 |
| 0 | *LOC108190590* | 0.969692 | 5.02E-05 |
| 0 | *hsp90ab1* | 0.554665 | 8.80E-05 |
| 0 | *eef1a1l1* | 0.385662 | 0.000138 |
| 0 | *pkp3a* | 0.755279 | 0.000384 |

| cluster | marker gene | avg_logFC | p_val_adj (<0.05) |
|---|---|---|---|
| 0 | *perp* | 0.767178 | 0.00053 |
| 0 | *tuba8l* | 0.69932 | 0.000539 |
| 0 | *cldne* | 0.672094 | 0.00054 |
| 0 | *gapdhs* | 0.653288 | 0.000545 |
| 0 | *ahnak* | 0.683779 | 0.000597 |
| 0 | *mcl1b* | 0.66905 | 0.000923 |
| 0 | *pabpc1a* | 0.430515 | 0.000996 |
| 0 | *cyp1a* | 0.742627 | 0.001284 |
| 0 | *zgc:162730* | 0.746051 | 0.001372 |
| 0 | *zgc:153665* | 0.73004 | 0.001454 |
| 0 | *si:ch211-195b11.3* | 0.813396 | 0.002542 |
| 0 | *ucp2* | 0.71683 | 0.003733 |
| 0 | *spint2* | 0.663588 | 0.003753 |
| 0 | *hspa8* | 0.352731 | 0.003822 |
| 0 | *rgs2* | 0.641279 | 0.00398 |
| 0 | *gsnb* | 0.471605 | 0.004024 |
| 0 | *krt15* | 1.155599 | 0.004226 |
| 0 | *cebpd* | 0.689705 | 0.004348 |
| 0 | *rbp4* | 0.660357 | 0.005953 |
| 0 | *capns1a* | 0.542324 | 0.006572 |
| 0 | *pnp5a* | 0.715041 | 0.007224 |
| 0 | *cd9b* | 0.747214 | 0.008575 |
| 0 | *ptmaa* | 0.499652 | 0.010898 |
| 0 | *ywhaz* | 0.539399 | 0.013014 |
| 0 | *higd1a* | 0.543086 | 0.015024 |
| 0 | *lye* | 0.829279 | 0.01534 |
| 0 | *gstm.3* | 0.808979 | 0.018662 |
| 0 | *nfkbiab* | 0.587787 | 0.019353 |
| 0 | *si:ch211-207n23.2* | 0.671449 | 0.023774 |
| 0 | *aqp3a* | 0.759478 | 0.024605 |
| 0 | *ccl25b* | 0.610525 | 0.030008 |
| 0 | *fkbp5* | 0.607871 | 0.042646 |
| 1 | *ubap2b* | 0.851111 | 7.06E-09 |
| 1 | *ND2* | 0.699785 | 2.43E-07 |
| 1 | *ND4* | 0.661873 | 9.49E-07 |
| 1 | *rsl24d1* | 0.660301 | 1.02E-06 |
| 1 | *vdac2* | 0.513986 | 1.67E-06 |
| 1 | *mt-co1* | 0.59156 | 2.25E-06 |
| 1 | *mt-nd1* | 0.700576 | 4.82E-06 |
| 1 | *ATP6* | 0.632829 | 6.49E-06 |
| 1 | *rpl34* | 0.39191 | 9.41E-06 |
| 1 | *si:ch211-199o1.2* | 0.69298 | 1.36E-05 |

| cluster | marker gene | avg_logFC | p_val_adj (<0.05) |
|---|---|---|---|
| 1 | *gapdh* | 0.474914 | 1.46E-05 |
| 1 | *rpl27a* | 0.536317 | 2.32E-05 |
| 1 | *mt-co2* | 0.558511 | 3.20E-05 |
| 1 | *riox1* | 0.772047 | 3.64E-05 |
| 1 | *gpx4a* | 0.613316 | 3.97E-05 |
| 1 | *si:dkey-151g10.6* | 0.418605 | 4.54E-05 |
| 1 | *rpl9* | 0.435873 | 7.31E-05 |
| 1 | *ND3* | 0.581373 | 7.87E-05 |
| 1 | *mkrn1* | 0.738568 | 8.41E-05 |
| 1 | *rpl35a* | 0.333371 | 8.58E-05 |
| 1 | *nsa2* | 0.613497 | 9.50E-05 |
| 1 | *hsp90aa1.1* | 0.512775 | 0.000125 |
| 1 | *si:ch211-131k2.3* | 0.52454 | 0.000145 |
| 1 | *rps27.1* | 0.372013 | 0.000167 |
| 1 | *XLOC-043851* | 0.585258 | 0.000296 |
| 1 | *rps21* | 0.414924 | 0.000297 |
| 1 | *rack1* | 0.323772 | 0.000347 |
| 1 | *rpl38* | 0.428235 | 0.00044 |
| 1 | *ddx21* | 0.559484 | 0.000483 |
| 1 | *tpt1* | 0.25937 | 0.000509 |
| 1 | *LOC101885209* | 0.45401 | 0.000515 |
| 1 | *rps14* | 0.372018 | 0.000595 |
| 1 | *mt-nd5* | 0.693903 | 0.000828 |
| 1 | *atp5if1b* | 0.481316 | 0.000831 |
| 1 | *hsc70* | 0.525304 | 0.001003 |
| 1 | *hspb1* | 0.569736 | 0.00111 |
| 1 | *rpl23* | 0.258052 | 0.001605 |
| 1 | *eef2l2* | 0.396717 | 0.001614 |
| 1 | *cast* | 0.530713 | 0.002765 |
| 1 | *COX3* | 0.43984 | 0.003425 |
| 1 | *rps11* | 0.303711 | 0.003921 |
| 1 | *rpl11* | 0.273629 | 0.004095 |
| 1 | *rps29* | 0.319585 | 0.004286 |
| 1 | *rps28* | 0.296043 | 0.004605 |
| 1 | *rplp2* | 0.37549 | 0.005792 |
| 1 | *rpl10a* | 0.267241 | 0.005916 |
| 1 | *si:dkey-51e6.1* | 0.489338 | 0.005973 |
| 1 | *rps13* | 0.26925 | 0.007281 |
| 1 | *pabpc4* | 0.408314 | 0.007572 |
| 1 | *btf3* | 0.30089 | 0.007798 |
| 1 | *calcoco2* | 0.45401 | 0.007854 |
| 1 | *rpl32* | 0.323661 | 0.009023 |

| cluster | marker gene | avg_logFC | p_val_adj (<0.05) |
|---|---|---|---|
| 1 | *NC-002333.17* | 0.727469 | 0.009119 |
| 1 | *rpl5b* | 0.293761 | 0.009652 |
| 1 | *rpl36* | 0.287416 | 0.010165 |
| 1 | *mt-cyb* | 0.434089 | 0.011919 |
| 1 | *ankrd39* | 0.338964 | 0.013806 |
| 1 | *psmb7* | 0.563705 | 0.014303 |
| 1 | *mlf1* | 0.505443 | 0.014779 |
| 1 | *rpl27* | 0.271985 | 0.015526 |
| 1 | *elocb* | 0.418875 | 0.015963 |
| 1 | *park7* | 0.587557 | 0.016573 |
| 1 | *rpl12* | 0.280456 | 0.020063 |
| 1 | *rpl26* | 0.253067 | 0.020954 |
| 1 | *si:ch211-214j24.10* | 0.442783 | 0.022261 |
| 1 | *rps26* | 0.267389 | 0.023725 |
| 1 | *dusp27* | 0.464821 | 0.023772 |
| 1 | *pinx1* | 0.511481 | 0.026918 |
| 1 | *rps27.2* | 0.277912 | 0.027535 |
| 1 | *rps23* | 0.281091 | 0.029009 |
| 1 | *psmc1b* | 0.617776 | 0.029329 |
| 1 | *hspd1* | 0.384414 | 0.035002 |
| 1 | *gyg1a* | 0.445903 | 0.039126 |
| 1 | *unc45b* | 0.442636 | 0.041465 |
| 1 | *pvalb7* | 0.592636 | 0.046203 |
| 1 | *rps25* | 0.257994 | 0.047109 |
| 2 | *ttn.1* | 0.857568 | 9.54E-11 |
| 2 | *si:ch73-367p23.2* | 0.856578 | 4.25E-10 |
| 2 | *mylz3* | 0.815089 | 6.75E-10 |
| 2 | *tnnt3b* | 0.654167 | 1.21E-09 |
| 2 | *tnni2a.3* | 1.188064 | 2.03E-09 |
| 2 | *mylpfa* | 0.623056 | 2.91E-09 |
| 2 | *myha* | 2.015147 | 5.81E-09 |
| 2 | *tnnc2* | 0.788208 | 2.22E-08 |
| 2 | *nme2b.2* | 0.700833 | 2.48E-08 |
| 2 | *actc1b* | 0.731837 | 3.00E-08 |
| 2 | *tpma* | 0.647884 | 3.52E-08 |
| 2 | *myl1* | 0.884009 | 5.91E-08 |
| 2 | *neb* | 0.711243 | 2.60E-07 |
| 2 | *pvalb2* | 0.610532 | 4.94E-07 |
| 2 | *ak1* | 0.560101 | 5.32E-07 |
| 2 | *actn3a* | 0.859317 | 5.61E-07 |
| 2 | *eno1a* | 0.724711 | 7.24E-07 |
| 2 | *myom1a* | 0.734073 | 1.67E-06 |

| cluster | marker gene | avg_logFC | p_val_adj (<0.05) |
|---|---|---|---|
| 2 | *ttn.2* | 0.696125 | 2.04E-06 |
| 2 | *gatm* | 0.664796 | 1.02E-05 |
| 2 | *ryr3* | 0.654347 | 1.31E-05 |
| 2 | *ryr1b* | 0.636466 | 2.38E-05 |
| 2 | *col1a2* | 0.772483 | 2.43E-05 |
| 2 | *lims2* | 0.665623 | 5.65E-05 |
| 2 | *col1a1b* | 0.8506 | 6.56E-05 |
| 2 | *myhz1.3* | 0.750464 | 9.00E-05 |
| 2 | *pvalb1* | 0.605546 | 0.000115 |
| 2 | *hhatla* | 0.660186 | 0.000167 |
| 2 | *acta1b* | 0.684778 | 0.000212 |
| 2 | *atp2a1l* | 0.407953 | 0.000375 |
| 2 | *spegb* | 0.566197 | 0.000409 |
| 2 | *smyd1a* | 0.603374 | 0.000553 |
| 2 | *XLOC-040108* | 0.448414 | 0.00057 |
| 2 | *wu:fj49a02* | 0.646856 | 0.000706 |
| 2 | *tent5c* | 0.541674 | 0.001489 |
| 2 | *tnni2a.4* | 0.515447 | 0.001747 |
| 2 | *slc25a4* | 0.555674 | 0.001787 |
| 2 | *myom2a* | 0.584397 | 0.001946 |
| 2 | *BX276101.1* | 0.562262 | 0.002286 |
| 2 | *col1a1a* | 0.685362 | 0.002544 |
| 2 | *frem3* | 0.42374 | 0.004415 |
| 2 | *CABZ01061524.1* | 0.582485 | 0.00461 |
| 2 | *klhl41b* | 0.538095 | 0.0052 |
| 2 | *ckmb* | 0.328592 | 0.005399 |
| 2 | *CABZ01072309.1* | 0.535945 | 0.006689 |
| 2 | *si:ch211-266g18.10* | 0.642426 | 0.008568 |
| 2 | *myom1b* | 0.656053 | 0.009682 |
| 2 | *tcap* | 0.541365 | 0.010362 |
| 2 | *si:ch211-255p10.3* | 0.550058 | 0.011476 |
| 2 | *fkbp1b* | 0.436393 | 0.011815 |
| 2 | *col6a3* | 0.374986 | 0.014445 |
| 2 | *col2a1b* | 0.642885 | 0.014656 |
| 2 | *ldb3a* | 0.50513 | 0.015785 |
| 2 | *XLOC-027924* | 0.503104 | 0.022395 |
| 2 | *serpinf1* | 0.450548 | 0.022599 |
| 2 | *lmnb1* | 0.307599 | 0.023106 |
| 2 | *mrpl27* | 0.350352 | 0.029784 |
| 2 | *igfn1.1* | 0.675762 | 0.032403 |
