## Supplementary figures and images for "Zebrafish *Danio rerio* trunk muscle structure and growth from a spatial transcriptomics perspective"

### experimental materials

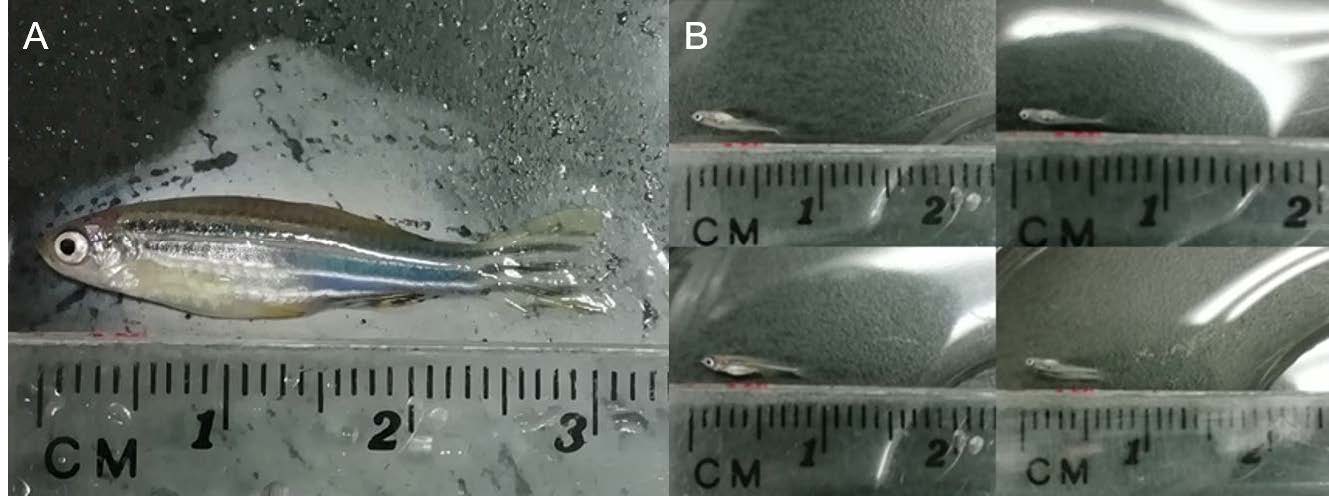

### Visium method

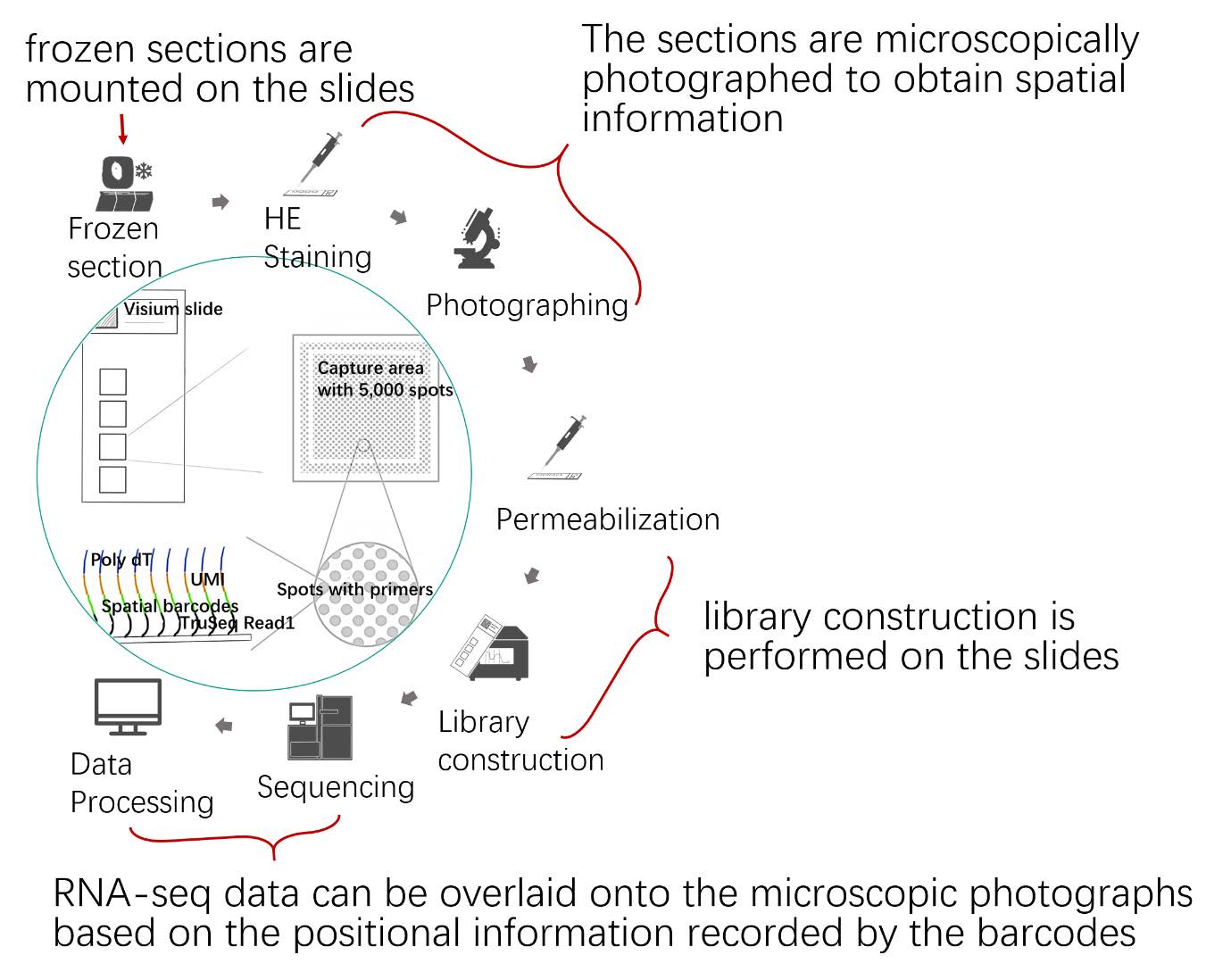
