## Supplementary material for "Zebrafish *Danio rerio* trunk muscle structure and growth from a spatial transcriptomics perspective": Permeabilization optimization and Quality check of cDNA libraries

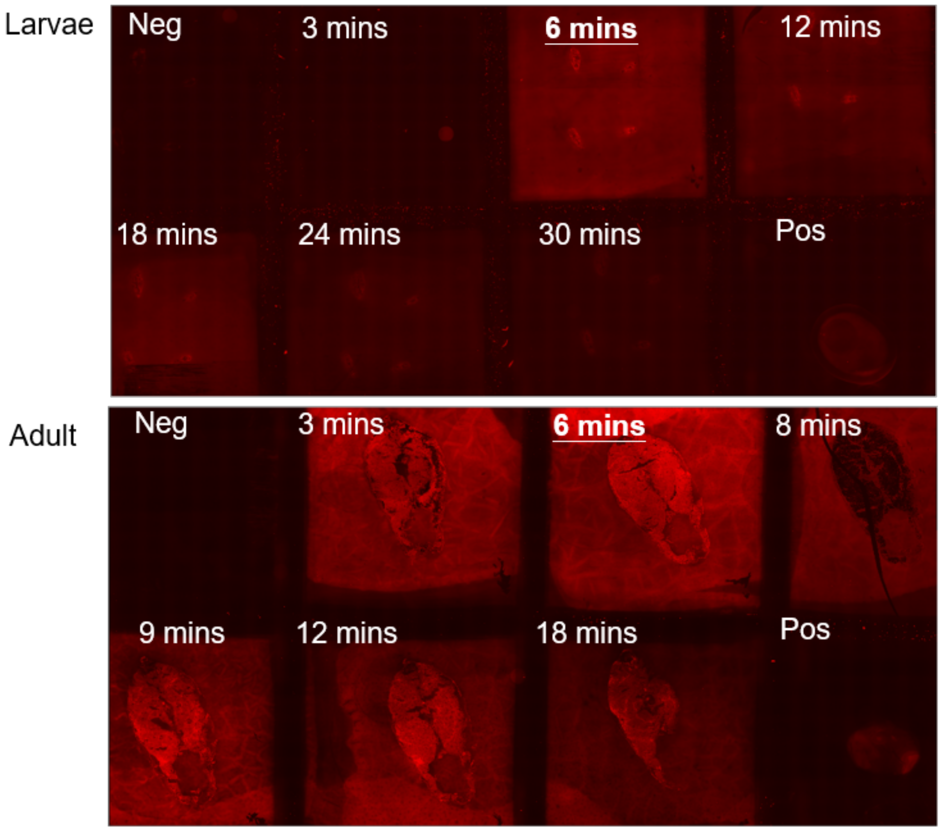


Supplementary 6 Tissue permeabilization optimization

Both Larvae and adult showed the clearest fluorescence at the permeabilization time of 6 min. Neg, negative control (no permeabilization). Pos, positive control (purified mRNA).

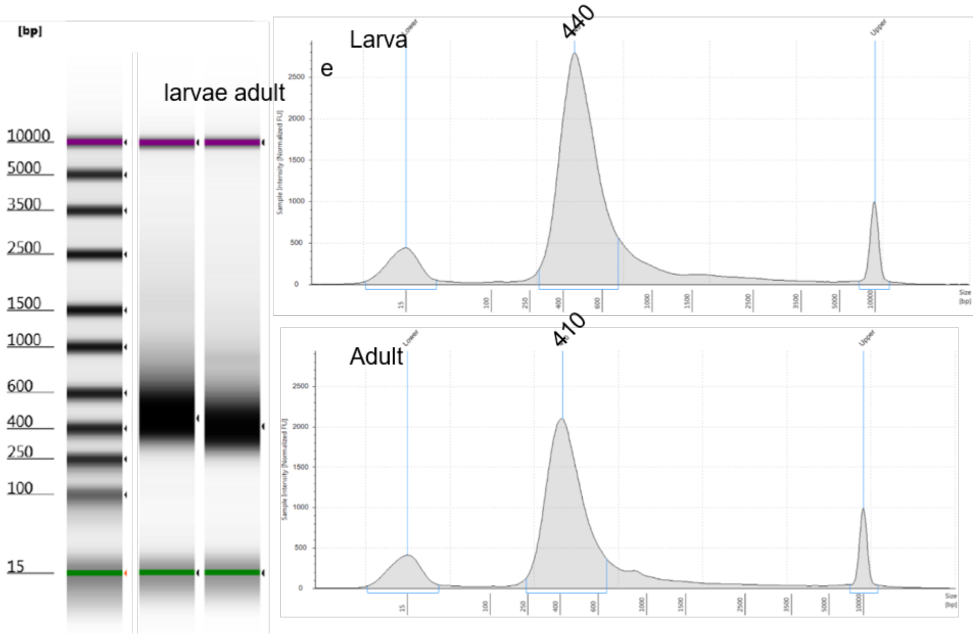


Supplementary 7 Quality check of cDNA libraries of larvae and adult

Electrophoresis pattern of constructed cDNA libraries by Tape Station.
